# Evidence for Functional Regulation of the KLHL3/WNK Pathway by O-GlcNAcylation

**DOI:** 10.1101/2025.02.27.640596

**Authors:** Jimin Hu, Duc T. Huynh, Denise E. Dunn, Jianli Wu, Cindy Manriquez-Rodriguez, Qianyi E. Zhang, Gabrielle A. Hirschkorn, George R. Georgiou, Tetsuya Hirata, Samuel A. Myers, Scott R. Floyd, Jen-Tsan Chi, Michael Boyce

## Abstract

The 42-member Kelch-like (KLHL) protein family are adaptors for ubiquitin E3 ligase complexes, governing the stability of a wide range of substrates. KLHL proteins are critical for maintaining proteostasis in a variety of tissues and are mutated in human diseases, including cancer, neurodegeneration, and familial hyperkalemic hypertension. However, the regulation of KLHL proteins remains incompletely understood. Previously, we reported that two KLHL family members, KEAP1 and gigaxonin, are regulated by O-linked β-*N*-acetylglucosamine (O-GlcNAc), an intracellular form of glycosylation. Interestingly, some ubiquitination targets of KEAP1 and gigaxonin are themselves also O-GlcNAcylated, suggesting that multi-level control by this post-translational modification may influence many KLHL pathways. To test this hypothesis, we examined KLHL3, which ubiquitinates with-no-lysine (WNK) kinases to modulate downstream ion channel activity. Our biochemical and glycoproteomic data demonstrate that human KLHL3 and all four WNK kinases (WNK1-4) are O-GlcNAcylated. Moreover, our results suggest that O-GlcNAcylation affects WNK4 function in both osmolarity control and ferroptosis, with potential implications ranging from blood pressure regulation to neuronal health and survival. This work demonstrates the functional regulation of the KLHL3/WNK axis by O-GlcNAcylation and supports a broader model of O-GlcNAc serving as a general regulator of KLHL signaling and proteostasis.

## Introduction

O-linked β-*N*-acetylglucosamine (O-GlcNAc) is a monosaccharide post-translational modification (PTM) on serine and threonine residues of cytoplasmic, nuclear, and mitochondrial proteins. In mammals, O-GlcNAc is added by O-GlcNAc transferase (OGT) and removed by O-GlcNAcase (OGA) (Yang, X. and Qian, K. 2017; Ong, Q., Han, W., et al. 2018; Hu, C.W., Wang, K., et al. 2024; Mayfield, J.M., Hitefield, N.L., et al. 2024; Udeshi, N.D., Hart, G.W., et al. 2024). O-GlcNAcylation governs numerous cellular processes, ranging from transcription and translation to metabolism and stress responses (Hart, G.W. 2014; Yang, X. and Qian, K. 2017; Ong, Q., Han, W., et al. 2018; Chatham, J.C., Zhang, J., et al. 2021), and dysregulation of O-GlcNAc signaling is implicated in many human diseases, including diabetes, neurodegenerative disorders, cancer, X-linked intellectual disability, and cardiovascular dysfunction (Chatham, J.C., Zhang, J., et al. 2021; Le Minh, G., Esquea, E.M., et al. 2023; Pratt, M.R. and Vocadlo, D.J. 2023; Mayfield, J.M., Hitefield, N.L., et al. 2024; Umapathi, P., Aggarwal, A., et al. 2024). Although O-GlcNAc was discovered decades ago, significant aspects of its function remain incompletely understood, including the key glycosylated substrates that mediate its downstream impact on many cell biological processes.

The Kelch-like (KLHL) protein family are adaptors for E3 ubiquitin ligase complexes and facilitate the ubiquitination and (usually) proteasomal degradation of substrate proteins (Genschik, P., Sumara, I., et al. 2013; Zhou, Y., Zhang, Q., et al. 2024). Humans have 42 KLHL proteins, all of which comprise an N-terminal BTB domain, a BACK domain, and C-terminal Kelch repeats (Dhanoa, B.S., Cogliati, T., et al. 2013; Zhou, Y., Zhang, Q., et al. 2024). The Kelch repeats recruit substrates for ubiquitination and are conserved from *Drosophila* to humans (Dhanoa, B.S., Cogliati, T., et al. 2013; Genschik, P., Sumara, I., et al. 2013; Zhou, Y., Zhang, Q., et al. 2024). Most KLHL proteins facilitate substrate degradation, with a few reported exceptions. For example, KLHL40 and KLHL41 stabilize proteins in the sarcomere through ubiquitination (Garg, A., O’Rourke, J., et al. 2014; Ramirez-Martinez, A., Cenik, B.K., et al. 2017). In contrast to their shared biochemical features, KLHL family members differ in their expression patterns and physiological functions and have been implicated in a range of disorders, including several types of cancer, hypertension, and neuromuscular disease (Dhanoa, B.S., Cogliati, T., et al. 2013; Fu, A.B., Xiang, S.F., et al. 2023; Zhou, Y., Zhang, Q., et al. 2024). Considerable progress has been made in understanding the functions of KLHL proteins, but how KLHL ubiquitination pathways are regulated under different physiological and pathological contexts remains largely unclear.

Previously, we reported that the KLHL proteins KEAP1 and gigaxonin are both O-GlcNAcylated, and we mapped O-GlcNAc sites on both proteins by mass spectrometry (MS) (Chen, P.H., Smith, T.J., et al. 2017; Chen, P.H., Hu, J., et al. 2020). We further found that site-specific O-GlcNAcylation regulates substrate degradation and protein-protein interactions for both proteins (Chen, P.H., Smith, T.J., et al. 2017; Chen, P.H., Hu, J., et al. 2020). Intriguingly, several KLHL ubiquitination targets are also themselves O-GlcNAcylated. For example, we and other have shown that the gigaxonin substrates vimentin and neurofilament-light are modified by O-GlcNAc to control their assembly states and functions (Dong, D.L., Xu, Z.S., et al. 1993; Slawson, C., Lakshmanan, T., et al. 2008; Chen, P.H., Smith, T.J., et al. 2017; Tarbet, H.J., Dolat, L., et al. 2018; Chen, P.H., Hu, J., et al. 2020; Huynh, D.T., Tsolova, K.N., et al. 2023). Other groups have reported the O-GlcNAcylation of NRF2, the canonical target of KEAP1-mediated ubiquitination and a master transcriptional regulator of redox stress responses (Costa, R.M., Dias, M.C., et al. 2024; Zhang, Y., Sun, C., et al. 2024; Yang, L., Tang, H., et al. 2025). Taken together, these results and the highly similar domain structures of KLHL proteins led us to hypothesize that O-GlcNAc may be a general, multi-level regulator of the KLHL family and its substrates, linking upstream signaling to downstream proteostasis.

As a next test of this hypothesis, we selected KLHL3, which influences ion homeostasis in the kidney and blood pressure control through regulated degradation of with-no-lysine (WNK) kinases (Ohta, A., Schumacher, F.R., et al. 2013; Shibata, S., Zhang, J., et al. 2013; Wakabayashi, M., Mori, T., et al. 2013). In response to stimulation by the vasoconstrictive hormone angiotensin II, protein kinase C phosphorylates KLHL3 at S433, which inhibits KLHL3 activity, reducing WNK ubiquitination and increasing WNK stability and kinase activity (Shibata, S., Arroyo, J.P., et al. 2014). WNKs then phosphorylate and activate the downstream kinases SPAK and OSR1, which in turn phosphorylate and activate ion transporters, such as the sodium-chloride cotransporter (NCC) (Hadchouel, J., Ellison, D.H., et al. 2016; Shekarabi, M., Zhang, J., et al. 2017; Ferdaus, M.Z. and McCormick, J.A. 2018; Uchida, S., Mori, T., et al. 2022). NCC hyperactivity leads to increased sodium and chloride reabsorption in the kidney, disrupting ion homeostasis and changing cellular osmolarity (Shekarabi, M., Zhang, J., et al. 2017; Uchida, S., Mori, T., et al. 2022). Therefore, the KLHL3/WNK pathway is critical for blood pressure regulation. Indeed, mutations in KLHL3, WNK1, WNK4, and the E3 ligase scaffold protein CUL3 cause familial hyperkalemic hypertension (FHHt), which is characterized by high blood pressure and high serum potassium (Wilson, F.H., Disse-Nicodeme, S., et al. 2001; Boyden, L.M., Choi, M., et al. 2012; Louis-Dit-Picard, H., Barc, J., et al. 2012; Maeoka, Y., Cornelius, R.J., et al. 2023). WNK signaling is critical in other tissues as well. For example, in the nervous system, WNK kinases regulate processes such as chloride sensing, GABA signaling, proliferation, and development, with pathological implications in the ischemic injury response, neuropathies, and brain tumors (Tang, B.L. 2016; Yang, T., Zhao, K., et al. 2017; Chen, W., Zebaze, L.N., et al. 2018; Murillo-de-Ozores, A.R., Chavez-Canales, M., et al. 2020; Izadifar, A., Courchet, J., et al. 2021; Bhuiyan, M.I.H., Young, C.B., et al. 2022; Shimizu, M. and Shibuya, H. 2022; Goldsmith, E.J. and Rodan, A.R. 2023; Zhang, H., Wang, Z., et al. 2024). Given its importance in tissue physiology and human disease, understanding the regulation of the KLHL3/WNK axis is a high priority.

Here, we report that both KLHL3 and WNK kinases are O-GlcNAcylated, and we provide evidence for functional effects of WNK4 O-GlcNAcylation in both osmosensation and ferroptosis. Though WNK kinases are well-known osmoregulators (Murillo-de-Ozores, A.R., Chavez-Canales, M., et al. 2020; Boyd-Shiwarski, C.R., Shiwarski, D.J., et al. 2022; Goldsmith, E.J. and Rodan, A.R. 2023), their involvement in ferroptosis is largely unexplored. Therefore, our results inform both established and emerging aspects of WNK biology. Taken together, our studies indicate a new role for O-GlcNAcylation in the KLHL3/WNK signaling pathway and support the larger hypothesis that O-GlcNAc regulates proteostasis in part through KLHL proteins and their substrates.

## Results

### KLHL3 is dynamically O-GlcNAcylated

To determine whether KLHL3 is O-GlcNAcylated, we expressed human KLHL3-myc-Flag in HEK293T cells treated with or without the small molecule OGA inhibitor Thiamet-G (TG) (Yuzwa, S.A., Macauley, M.S., et al. 2008). Immunoprecipitation and Western blot (IP/WB) analysis demonstrated that KLHL3 is O-GlcNAcylated (Figure 1A). We verified this result with hexosaminidase treatment, which cleaves O-GlcNAc from proteins (Magnelli, P., Bielik, A., et al. 2012), confirming the specificity of the O-GlcNAc signal on KLHL3 (Figure 1B). Next, we expressed KLHL3-myc-6xHis in HEK293T cells, purified the protein to homogeneity, and performed MS analysis to identify specific O-GlcNAc sites, as we have done previously (Myers, S.A., Daou, S., et al. 2013; Myers, S.A., Peddada, S., et al. 2016; Chen, P.H., Smith, T.J., et al. 2017; Cox, N.J., Unlu, G., et al. 2018; Chen, P.H., Hu, J., et al. 2020; Bisnett, B.J., Condon, B.M., et al. 2021; Huynh, D.T., Tsolova, K.N., et al. 2023). MS results demonstrated that T264 is O-GlcNAcylated (Figures 1C and S1A). Interestingly, this residue has also been reported as a phosphorylation site in a prior MS study (Shibata, S., Arroyo, J.P., et al. 2014), suggesting potential PTM crosstalk between phosphorylation and O-GlcNAcylation on KLHL3.

**Figure 1.**
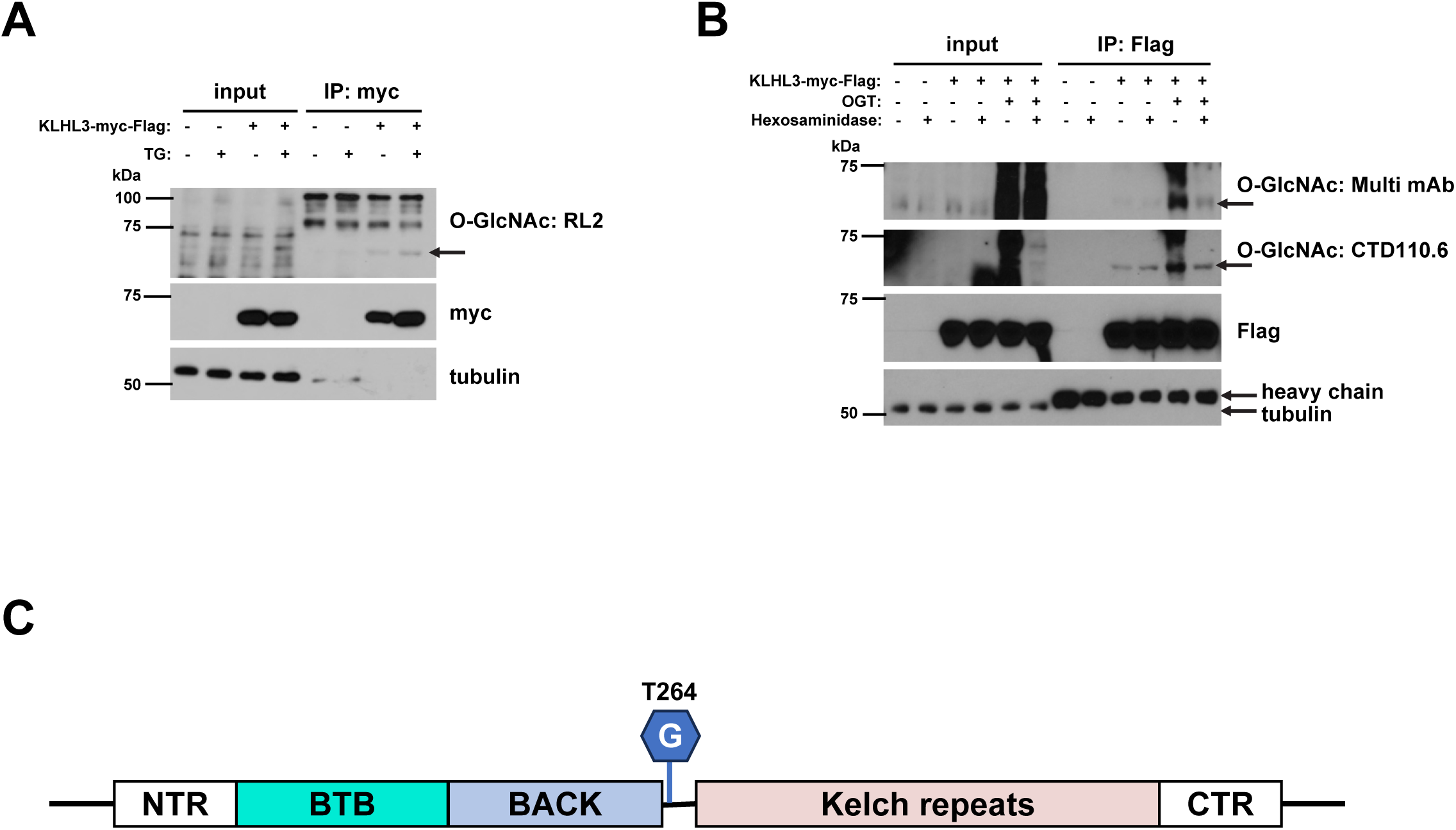
KLHL3 is dynamically O-GlcNAcylated. (A) HEK293T cells were transfected with KLHL3-myc-Flag for 24 hours and then treated with DMSO (vehicle) or 50 µM Thiamet-G (TG) for an additional 24 hours. Lysates were analyzed by IP/WB. n=3. (B) HEK293T cells were co-transfected with KLHL3-myc-Flag and empty vector or OGT-myc-6xHis for 48 hours. Lysates were treated with water (vehicle) or hexosaminidase for 1 hour at 37 °C and analyzed by IP/WB. n=3 for CTD110.6, Flag, and tubulin, n=2 for Multi mAb. (C) Schematic showing the position of T264, the O-GlcNAc site on KLHL3 identified by MS.

### WNK kinases are dynamically O-GlcNAcylated

We previously showed that O-GlcNAcylation of the KLHL proteins KEAP1 and gigaxonin influences their respective protein-protein interactions (Chen, P.H., Smith, T.J., et al. 2017; Chen, P.H., Hu, J., et al. 2020). Therefore, we sought to determine whether O-GlcNAcylation of KLHL3 affects its interaction with its best-studied substrate, WNK4 (Ohta, A., Schumacher, F.R., et al. 2013; Shibata, S., Zhang, J., et al. 2013; Wakabayashi, M., Mori, T., et al. 2013; Shibata, S., Arroyo, J.P., et al. 2014; Ishizawa, K., Wang, Q., et al. 2019; Murillo-de-Ozores, A.R., Rodriguez-Gama, A., et al. 2021). Initial co-IPs between KLHL3 and WNK4 showed no obvious O-GlcNAc-dependent changes in their interaction (Figure 2A). However, in this experiment, we unexpectedly observed a robust O-GlcNAc signal on WNK4, providing the first evidence that it might be glycosylated (Figure 2A). Indeed, follow-up IP/WB demonstrated that WNK4 is dynamically O-GlcNAcylated (Figure 2B). Treatment with the OGT inhibitor Ac_4_5SGlcNAc (5SG) (Gloster, T.M., Zandberg, W.F., et al. 2011) dramatically reduced the O-GlcNAc signal on WNK4, whereas TG raised it, confirming specificity (Figure 2B). We further showed that WNKs 1 through 3 are also O-GlcNAcylated (Figure 2C-E), suggesting that glycosylation may be a general regulator of both KLHL3 and all four WNK kinases.

**Figure 2.**
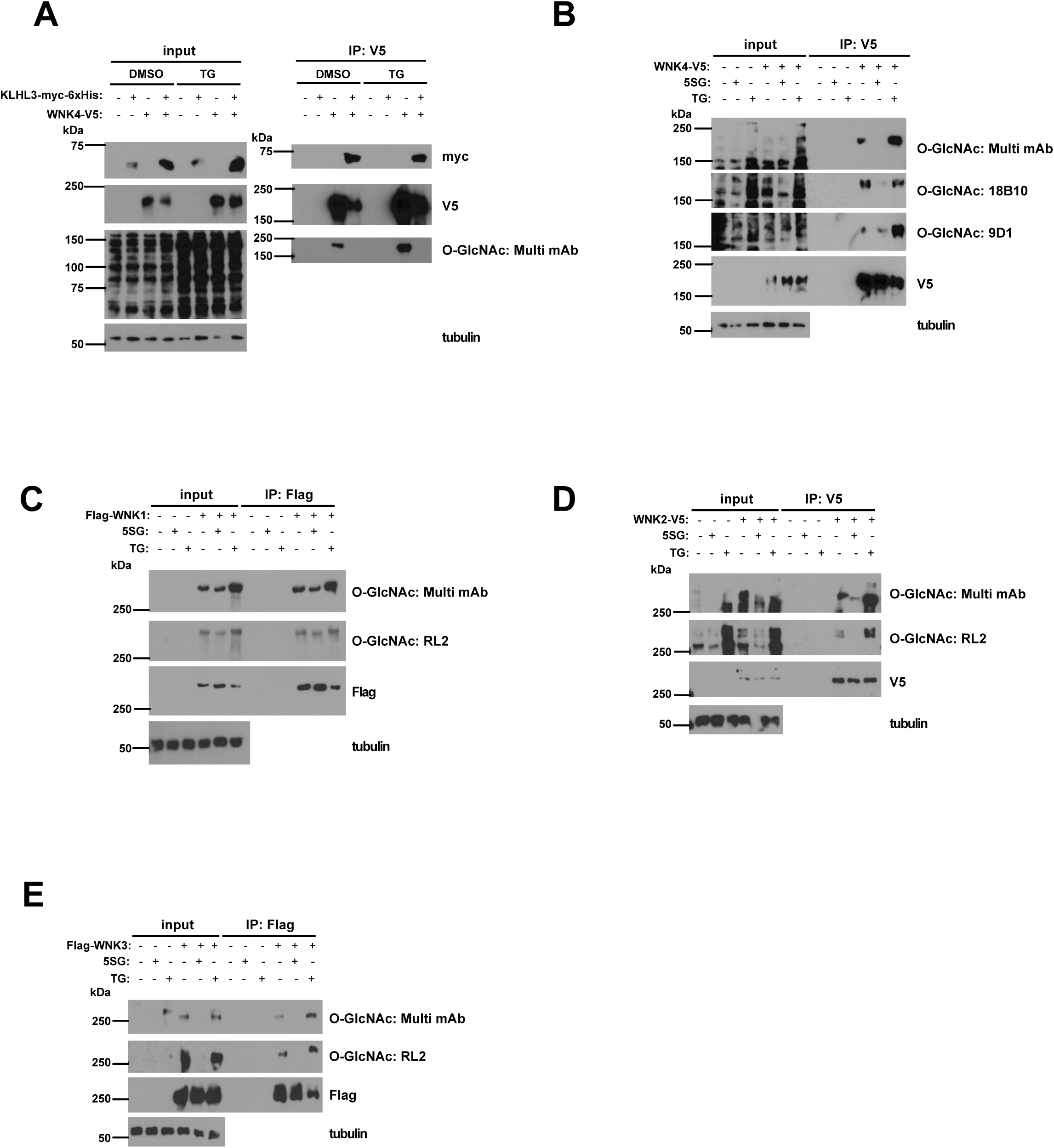
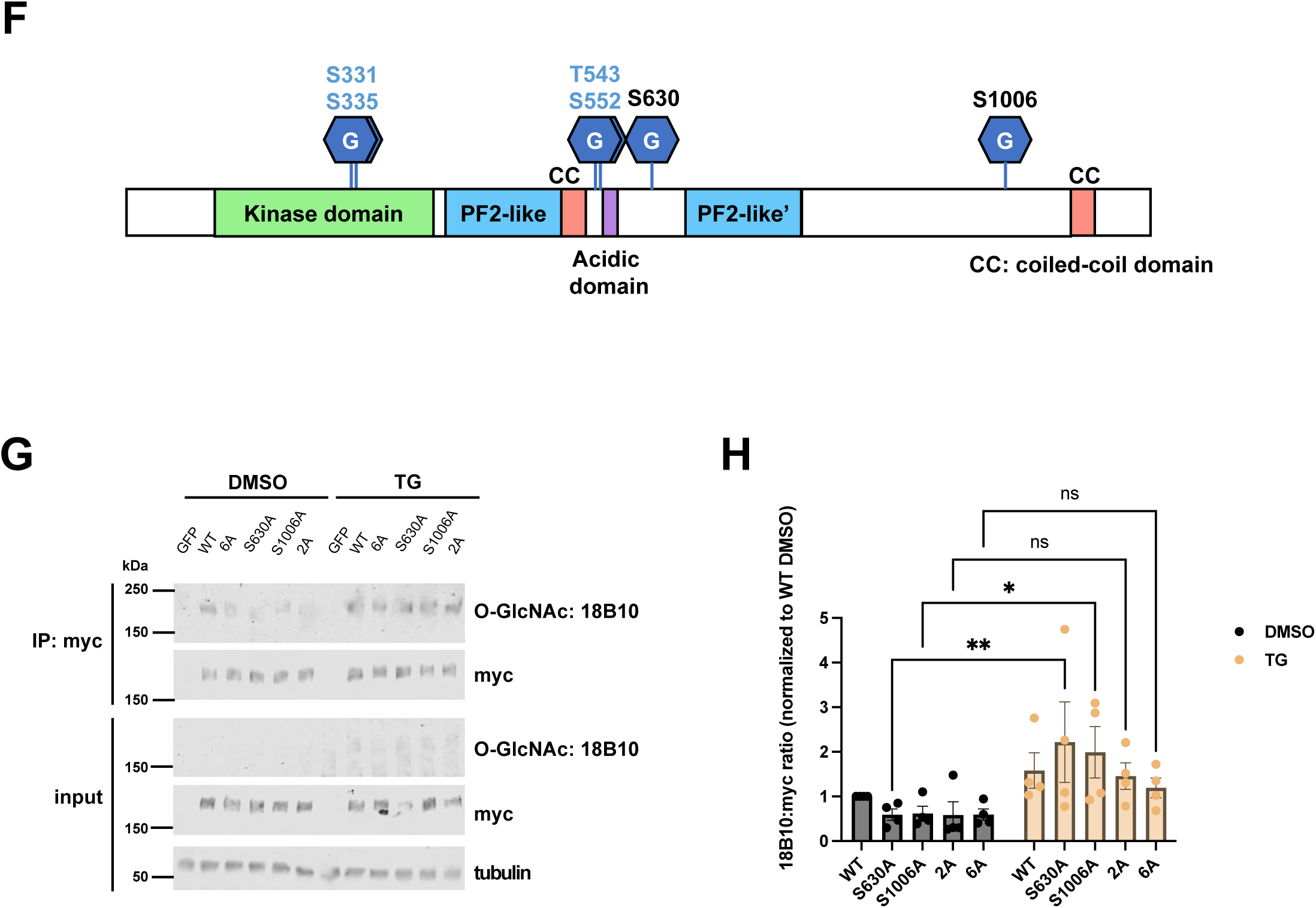
WNK kinases are dynamically O-GlcNAcylated. (A) HEK293T cells were transfected with KLHL3-myc-6xHis and/or WNK4-V5 for 24 hours and then treated with DMSO or 50 µM TG for an additional 24 hours. Lysates were analyzed by IP/WB. n=1. WNK4 levels are lower in cells co-transfected with KLHL3 because KLHL3 mediates WNK4 ubiquitination and degradation. (B) HEK293T cells were transfected with WNK4-V5 for 24 hours and then treated with DMSO, 50 µM Ac_4_5SGlcNAc (5SG), or 50 µM TG for an additional 24 hours. Lysates were analyzed by IP/WB. n=3. (C-E) HEK293T cells were transfected with each WNK plasmid as indicated for 24 hours and then treated with DMSO, 50 µM 5SG, or 50 µM TG for an additional 24 hours. Lysates were analyzed by IP/WB. n=3. (F) Schematic of WNK4 O-GlcNAc sites identified by MS. S630 and S1006 are ETD-confirmed sites, while the other four sites are candidates based on HCD MS-identified glycopeptides. (G) HEK293T cells were transfected with myc-6xHis-tagged WT or mutant WNK4 for 24 hours, then treated with DMSO or 50 µM TG for an additional 24 hours. Lysates were analyzed by IP/fluorescent WB. n=4. (H) Quantitation of WB data in (G). All data are displayed as mean ± SEM. Data were analyzed by two-way ANOVA followed by Tukey’s multiple comparisons test. n=4. *, *p* ≤ 0.05; **, *p* ≤ 0.01; ns, not significant.

As a first step toward elucidating the biochemical and cell biological effects of O-GlcNAcylation in the KLHL3/WNK axis, we focused on WNK4. We performed O-GlcNAc site-mapping on human WNK4 and identified two definitively identified sites and four additional candidate sites (which could not be unambiguously assigned from the fragmentation data) (Figures 2F and S1B). Interestingly, the confirmed S1006 O-GlcNAc site was also observed as a phosphorylation site in our MS datasets, suggesting the potential for PTM crosstalk on this residue. Of the four candidate O-GlcNAc sites that could not be unambiguously assigned, S335 is a reported autophosphorylation site, which is important for WNK4 activation under certain conditions, such as low intracellular chloride (Bazua-Valenti, S., Chavez-Canales, M., et al. 2015; Murillo-de-Ozores, A.R., Rodriguez-Gama, A., et al. 2021).

To determine whether O-GlcNAcylation is inducible at the residues we identified by MS, we created unglycosylatable alanine point-mutants for each of the confirmed WNK4 O-GlcNAc sites (S630 and S1006), a “2A” mutant for both confirmed sites (S630A/S1006A), and a “6A” mutant for all six sites, including the four candidates. We then quantified the O-GlcNAc levels of wild type (WT) WNK4 and selected mutants under control (DMSO) or TG-treated conditions (Figure 2G-H). The S630A and S1006A single mutants both exhibited increased O-GlcNAcylation in response to TG treatment, relative to vehicle only (Figure 2H). However, both the 2A and the 6A mutants showed no statistically significant increase in O-GlcNAcylation under TG treatment (Figure 2H). These observations suggest that TG-induced O-GlcNAcylation may occur at S630 and S1006 independently of each other, and these two sites may be responsible, at least in part, for the notable increase in O-GlcNAcylation observed in WT WNK4 upon TG treatment (Figure 2B).

### Evidence for WNK4 as an O-GlcNAc-regulated osmolarity sensor

To assess the potential function of WNK glycosylation, we tested whether upstream stimuli affect WNK4 O-GlcNAcylation. WNK1 plays a well-established and important role in osmoregulation (Murillo-de-Ozores, A.R., Chavez-Canales, M., et al. 2020; Goldsmith, E.J. and Rodan, A.R. 2023), and osmotic stress impacts WNK4 phosphorylation (Maruyama, J., Kobayashi, Y., et al. 2016), suggesting that WNK PTMs mediate osmoregulation. To determine whether WNK4 O-GlcNAcylation participates in osmoregulation, we subjected HEK293T cells expressing WNK4 WT or 6A to different osmolarities and used IP/WB to quantify WNK4 O-GlcNAc levels (Figure 3A-B). WT WNK4 O-GlcNAcylation was significantly increased under hypertonic conditions, relative to the hypotonic condition, whereas the 6A mutant exhibited no significant change (Figure 3A-B). We also examined phosphorylation at S575, the WNK4 phosphosite reported to be modulated by osmotic stress (Maruyama, J., Kobayashi, Y., et al. 2016) (Figure S2). We did not detect the osmolarity-dependent changes in S575 phosphorylation reported previously for WT WNK4 (Maruyama, J., Kobayashi, Y., et al. 2016), perhaps due to differences in detection methods or experimental systems (Figure S2). However, under hypertonic conditions, we did observe a difference in phospho-S575 signal on WT versus 6A WNK4 (Figure S2), suggesting potential crosstalk between WNK4 phosphorylation and O-GlcNAcylation under osmotic stress.

**Figure 3.**
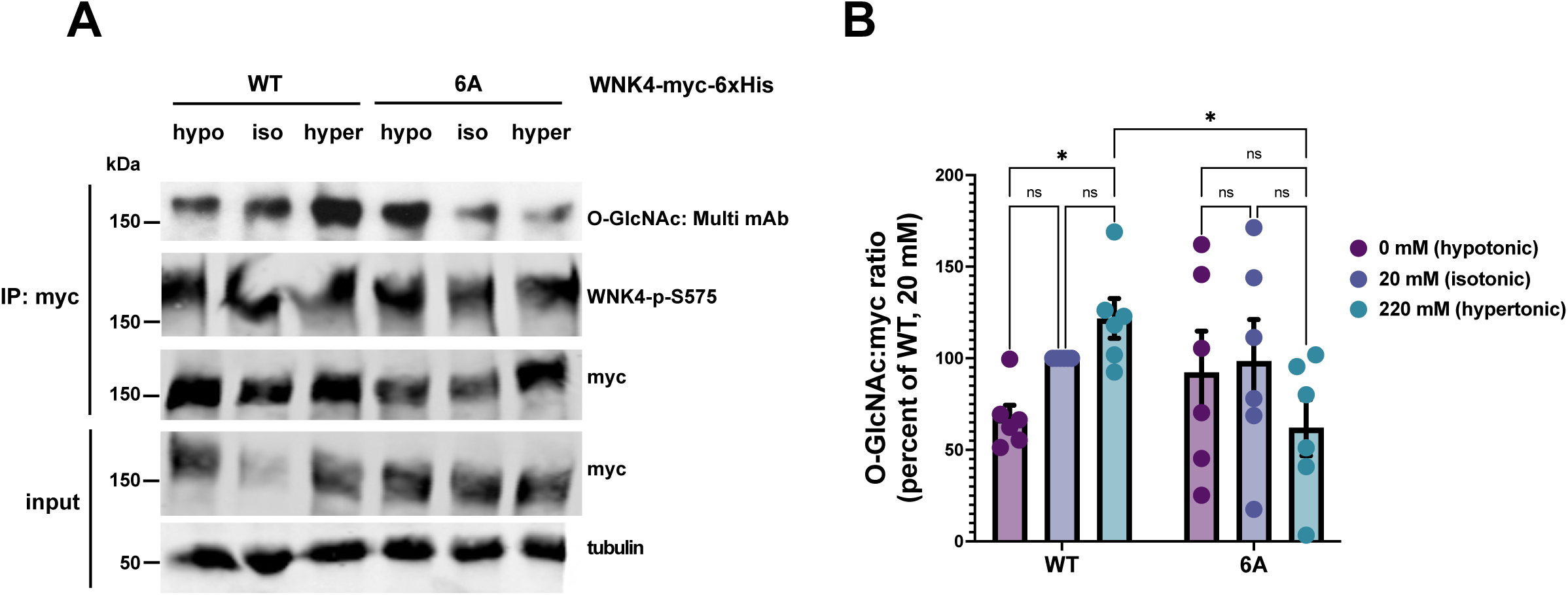
WNK4 O-GlcNAcylation is regulated by osmolarity. (A) HEK293T cells were transfected with WT or 6A WNK4-myc-6xHis for 24 hours and then incubated with hypotonic, isotonic, or hypertonic buffer, as indicated, for 30 minutes. Lysates were analyzed by IP/WB. All blots depicted were detected by fluorescence except the Multi mAb blot (enhanced chemiluminescence). n=6. (B) Fluorescent WB results from (A) were analyzed by two-way ANOVA with Tukey’s multiple comparisons test. All data are displayed as mean ± SEM. n=6. *, *p* ≤ 0.05; ns, not significant.

WNK4 has a C-terminal disordered region that is homologous to the WNK1 domain that mediates phase separation upon osmotic stress (Murillo-de-Ozores, A.R., Rodriguez-Gama, A., et al. 2021; Boyd-Shiwarski, C.R., Shiwarski, D.J., et al. 2022). Prior literature also indicates that hypertonicity changes WNK4 cellular localization, resulting in punctate morphology that resembles phase-separated WNK1 (Shaharabany, M., Holtzman, E.J., et al. 2008). Therefore, we investigated the potential connections among osmolarity and WNK4 O-GlcNAcylation and relocalization. For these studies, we selected 786-O human renal adenocarcinoma cells because they are a well-established model for kidney cell biology and are amenable to imaging studies (Satcher, R.L., Pan, T., et al. 2014; Xu, Z.Q., Zhang, L., et al. 2015; Zhao, C.X., Luo, C.L., et al. 2015; Brodaczewska, K.K., Szczylik, C., et al. 2016; Ishibashi, K., Koguchi, T., et al. 2018; Zou, Y., Palte, M.J., et al. 2019; Sevilla-Montero, J., Bienes-Martinez, R., et al. 2020; Zhang, X., Sun, Y., et al. 2023). We treated 786-O cells with OGT or OGA inhibitors, subjected them to different osmolarities, and visualized endogenous WNK4 by immunofluorescence (IF) (Figure 4A). In vehicle-treated cells, WNK4 was more diffuse and less punctate under hypotonic conditions, compared to isotonic or hypertonic (Figure 4A). However, either inhibition of OGT by OSMI-1 (Ortiz-Meoz, R.F., Jiang, J., et al. 2015) or of OGA by TG abolished these osmolarity-dependent differences (Figures 4A and S3). Average per-cell WNK4 fluorescence area dramatically decreased as osmolarity increased in vehicle-treated cells, but these effects were abrogated by OGT or OGA inhibition (Figure 4B). Notably, the punctate distribution of WNK4 under hypertonic conditions resembles the relocalization and phase separation behavior observed in prior WNK studies (Shaharabany, M., Holtzman, E.J., et al. 2008; Boyd-Shiwarski, C.R., Shiwarski, D.J., et al. 2022). Together, these results suggest that WNK4 is a sensor for osmotic stress in a kidney cell model and responds to osmolarity changes via relocalization and perhaps phase separation. Moreover, WNK4 O-GlcNAcylation is responsive to osmolarity changes, and O-GlcNAc cycling is required for WNK relocalization upon osmotic stress.

**Figure 4.**
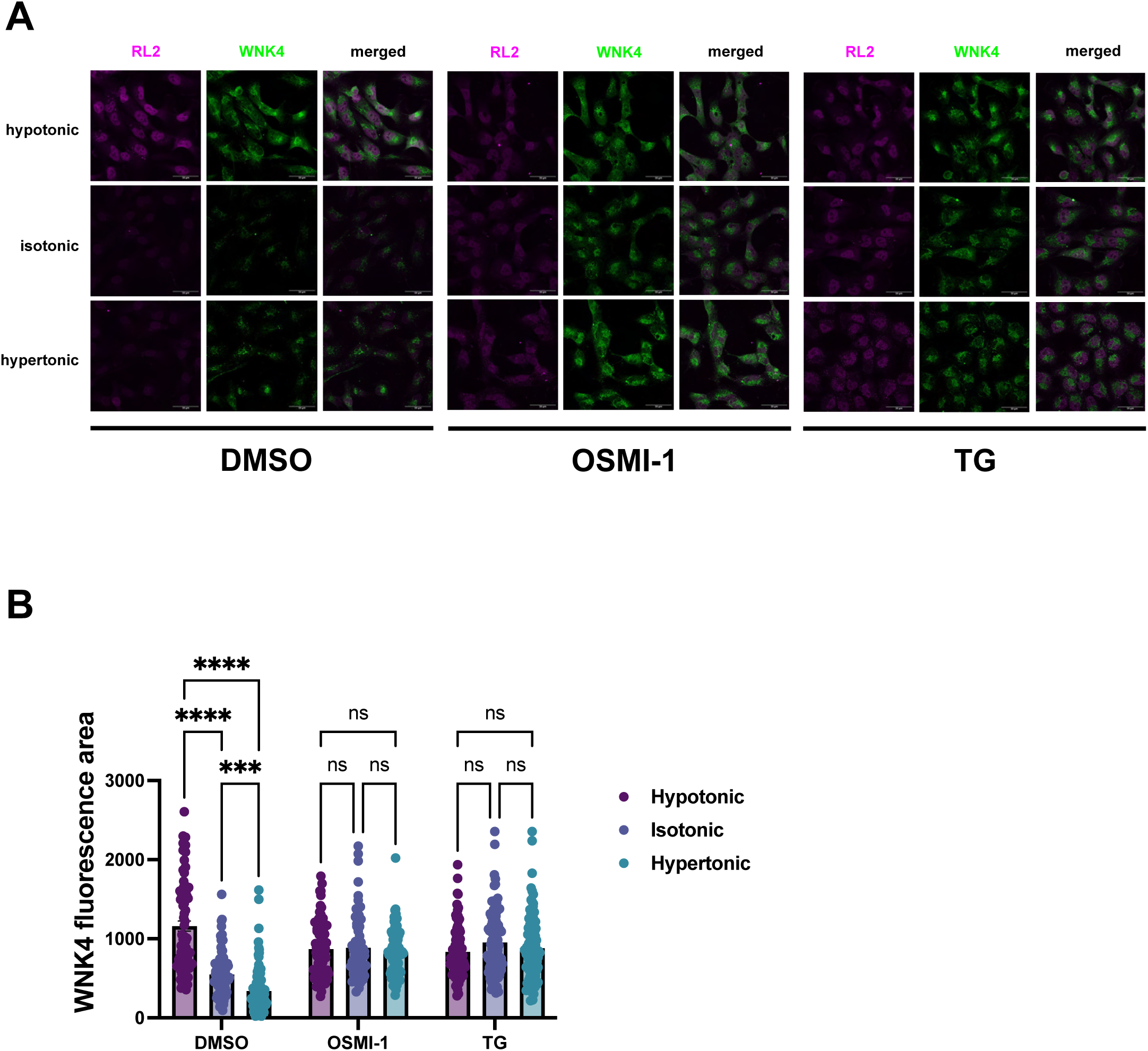
WNK4 osmosensation is O-GlcNAc-dependent. (A) 786-O cells were treated with DMSO, 50 µM TG, or 50 µM OSMI-1 for 30 minutes, incubated with hypotonic, isotonic, or hypertonic buffer supplemented with inhibitors for another 30 minutes (total inhibitor treatment time, 1 hour), and analyzed by IF. n=3. (B) WNK4 fluorescence area in individual cells was calculated from the experiment in (A). At least 20 cells were analyzed per condition in each biological replicate, and the graph represents data from n=3 biological replicates. All data are displayed as mean ± SEM. Data were analyzed by two-way ANOVA with Tukey’s multiple comparisons test. ***, *p* ≤ 0.001; ****, *p* ≤ 0.0001; ns, not significant.

### Evidence for WNK4 O-GlcNAcylation and activity in mediating ferroptosis

To examine the potential functional significance of WNK O-GlcNAcylation in another context, we turned to ferroptosis. Ferroptosis is a form of non-apoptotic cell death dependent on iron metabolism (Jiang, X., Stockwell, B.R., et al. 2021; Liang, D., Minikes, A.M., et al. 2022; Zhang, H., Zhang, J., et al. 2023). A key feature of ferroptosis is an increase in reactive oxygen species, which impairs cell viability in multiple ways, including via lipid and membrane damage (Jiang, X., Stockwell, B.R., et al. 2021; Liang, D., Minikes, A.M., et al. 2022; Zhang, H., Zhang, J., et al. 2023). Ferroptosis has been observed in many cell types in the nervous system, including neurons, microglia, astrocytes, and oligodendrocytes (Li, Y., Xiao, D., et al. 2022), with implications in neurodegenerative diseases such as Alzheimer’s Disease (AD), Parkinson’s Disease, and Huntington’s Disease (Stockwell, B.R., Friedmann Angeli, J.P., et al. 2017; Jakaria, M., Belaidi, A.A., et al. 2021; Li, Y., Xiao, D., et al. 2022; Wang, Z.L., Yuan, L., et al. 2022; Du, L., Wu, Y., et al. 2023; Fei, Y. and Ding, Y. 2024).

Recently, O-GlcNAc signaling has been implicated in multiple processes related to ferroptosis, such as oxidative stress regulation and lipid and iron metabolism (Rao, X., Duan, X., et al. 2015; Chen, P.H., Smith, T.J., et al. 2017; Chen, P.H., Chi, J.T., et al. 2018; Sodi, V.L., Bacigalupa, Z.A., et al. 2018; Chen, Y., Zhu, G., et al. 2019; Zhu, G., Murshed, A., et al. 2021; Yang, Z., Wei, X., et al. 2023; Zhang, H., Zhang, J., et al. 2023; Yang, L., Tang, H., et al. 2025). The role of WNK kinases in cell growth and survival has also been explored in different contexts, including, but not limited to, the nervous system (Park, H.W., Kim, J.Y., et al. 2011; Tu, S.W., Bugde, A., et al. 2011; Tang, B.L. 2016; Sanchez-Fdez, A., Matilla-Almazan, S., et al. 2023). Of particular interest, a recent study reported that WNK activity promotes ferroptotic cell death and white matter lesions (Xu, X., Sun, Y., et al. 2024). Taken together, these data suggest the hypothesis that WNK4 O-GlcNAcylation may play a role in mediating ferroptosis. Therefore, we investigated this notion next.

We first tested whether WNK4 participates in ferroptotic cell death. We treated control and WNK4 knockdown 786-O cells (Figure S4) with erastin, an inhibitor of the cystine-glutamate antiporter system Xc^-^, which causes ferroptosis by limiting cystine supply and disrupting the function of the phospholipid peroxidase GPX4, resulting in the accumulation of lipid peroxides that lead to ferroptosis (Dolma, S., Lessnick, S.L., et al. 2003; Dixon, S.J., Lemberg, K.M., et al. 2012; Dixon, S.J., Patel, D.N., et al. 2014; Lu, B., Chen, X.B., et al. 2017; Jiang, X., Stockwell, B.R., et al. 2021) (Figure 5A). Partial knockdown of endogenous WNK4 was sufficient to protect cells from ferroptosis, indicating that WNK4 participates in ferroptotic death in 786-O cells (Figures S4 and 5A).

**Figure 5.**
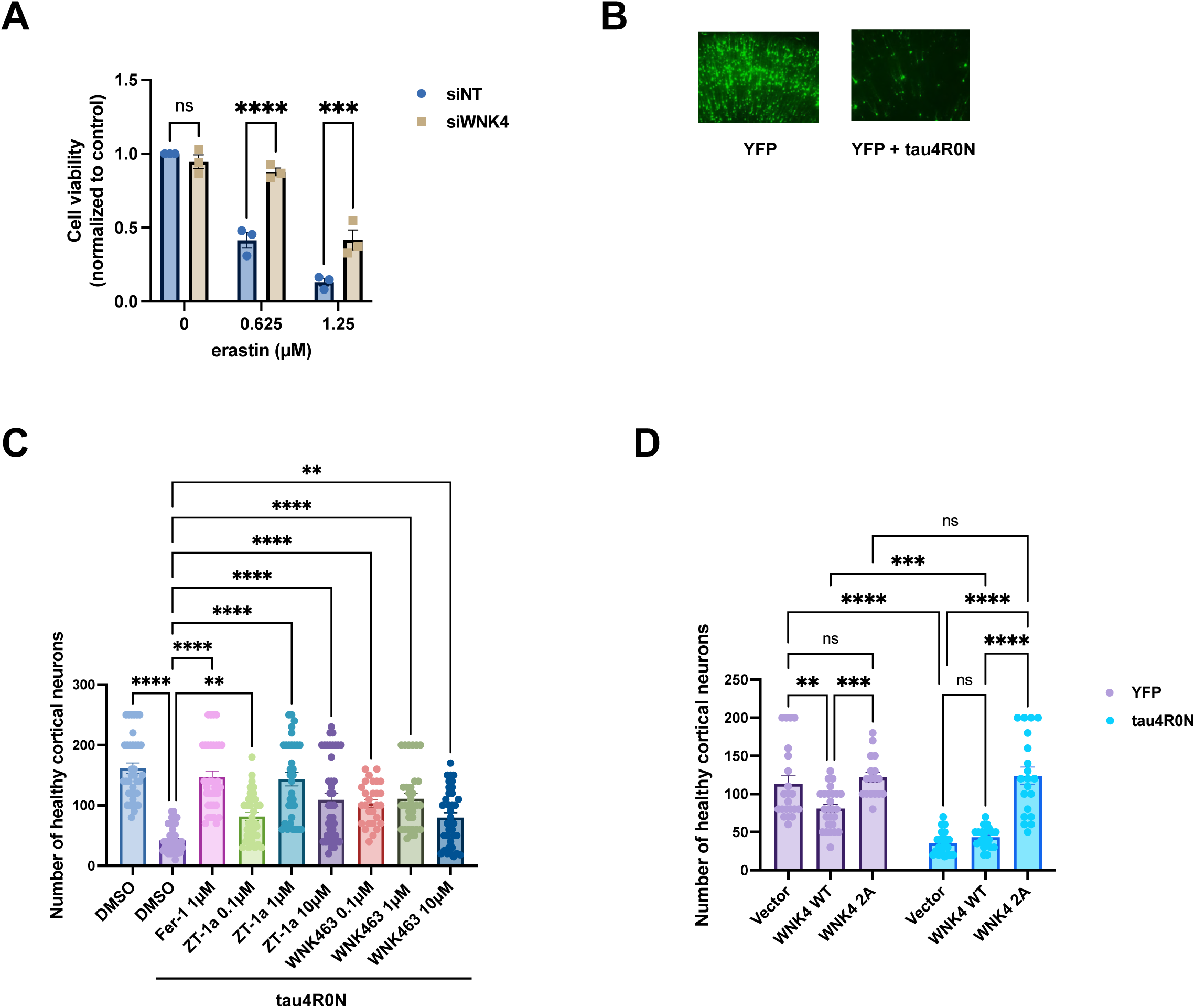
WNK4 activity and O-GlcNAcylation mediate ferroptosis. (A) 786-O cells were transfected with a non-targeting control siRNA (siNT) or a pooled siRNA against WNK4 (siWNK4) and treated with erastin doses as indicated for 24 hours. Cell viability was measured by the CellTiter-Glo assay. n=3, with three technical replicates per biological replicate. Data were normalized to the average of three technical replicates of siNT 0 µM erastin for each biological replicate, and the average of normalized values for three technical replicates of each condition was used as a data entry in the statistical analysis. Data were analyzed by two-way ANOVA, followed by Tukey’s multiple comparisons test. ****, *p* ≤ 0.0001; ns, not significant. (B) OBSCs were transfected with YFP only (left) or YFP + tau4R0N (right) for 72 hours and analyzed by fluorescence microscopy. (C) OBSCs were sectioned and placed in regular maintenance medium containing DMSO (vehicle), ferrostatin-1 (Fer-1), ZT-1a or WNK463, as indicated, immediately transfected with YFP only or YFP + tau4R0N, and analyzed by fluorescence microscopy after 72 hours. For each condition, 28-42 brain slices were included for quantitation, and the number of healthy cortical neurons in each brain slice was counted and plotted. Data were analyzed by one-way ANOVA followed by Dunnett’s multiple comparisons test. n=5. **, *p* ≤ 0.01; ****, *p* ≤ 0.0001. (D) OBSCs were transfected with YFP only (left) or YFP + tau4R0N (right) plus empty vector or WT or 2A WNK4 for 72 hours and were analyzed by fluorescence microscopy. For each condition, 17-26 brain slices were included for quantitation, and the number of healthy cortical neurons in each brain slice was counted and plotted. Data were analyzed by two-way ANOVA followed by Tukey’s multiple comparisons test. n=3. **, *p* ≤ 0.01; ***, *p* ≤ 0.001; ****, *p* ≤ 0.0001; ns, not significant. Throughout, all data are displayed as mean ± SEM.

To determine whether our WNK4/ferroptosis results extend to a more pathophysiologically relevant context, we took advantage of an established rat organotypic brain slice culture (OBSC) model of neurodegeneration that we have used previously (Braithwaite, S.P., Schmid, R.S., et al. 2010; Hoffstrom, B.G., Kaplan, A., et al. 2010; Reinhart, P.H., Kaltenbach, L.S., et al. 2011; Skouta, R., Dixon, S.J., et al. 2014; Van Kanegan, M.J., Dunn, D.E., et al. 2016; Venkatesh, D., O’Brien, N.A., et al. 2020; Chi, J.T., Lin, C.C., et al. unpublished data). In this version of the model, biolistic transfection of yellow fluorescent protein (YFP) alone or YFP plus a mutant form of the tau protein (tau4R0N) (Van Kanegan, M.J., Dunn, D.E., et al. 2016) into OBSCs (Figure 5B) causes tau-dependent cortical neuron death, which can be suppressed by the ferroptosis inhibitor ferrostatin-1 (Fer-1). Therefore, this OBSC system provides a tractable model of the tau-dependent, ferroptotic cell death component of neurodegeneration in AD. Although WNK4 has mostly been studied in the kidney, it is also expressed in the brain (Murillo-de-Ozores, A.R., Rodriguez-Gama, A., et al. 2021), and WNK signaling in general is known to influence neuronal survival (Tang, B.L. 2016; Chen, W., Zebaze, L.N., et al. 2018; Izadifar, A., Courchet, J., et al. 2021; Shimizu, M. and Shibuya, H. 2022), motivating these experiments.

To investigate the effects of the WNK/SPAK/OSR1 pathway on ferroptosis, we treated the OBSCs with the SPAK inhibitor ZT-1a (Zhang, J., Bhuiyan, M.I.H., et al. 2020) or the pan-WNK inhibitor WNK463 (Yamada, K., Park, H.M., et al. 2016), transfected with tau4R0N (or control), and quantified neuronal viability (Figure 5C). Both inhibitors increased cortical neuron viability in OBSCs, suggesting that the WNK/SPAK/OSR1 pathway sensitizes neurons to tau4R0N-induced ferroptosis in this context (Figure 5C). To extend these findings to another pathophysiologically relevant stimulus, we examined the ferroptosis caused by oxygen-glucose deprivation, a previously validated model for stroke in the OBSC system (Wang, J.K.T., Portbury, S., et al. 2006; Dunn, D.E., He, D.N., et al. 2011; Van Kanegan, M.J., Dunn, D.E., et al. 2016). Similar to tau-induced cell death, ZT-1a or WNK463 treatment protected against ferroptotic neuronal death, though the impact of WNK463 was more variable in this case (Figure S5). Finally, we examined the potential role of WNK4 O-GlcNAcylation in tau-induced neuronal death. Transfection of WT WNK4 decreased neuronal viability under YFP-only (control) conditions and had no significant impact on tau4R0N-induced cell death (Figure 5D). By contrast, expression of the WNK4 2A mutant did not impact neuronal viability under control conditions but significantly suppressed tau-induced cell death (Figure 5D). Taken together, our results demonstrate a role for WNK kinase activity and validated WNK4 O-GlcNAcylation sites in tau-induced ferroptotic cell death of cortical neurons in the OBSC model.

## Discussion

Here, we report the site-specific O-GlcNAcylation of KLHL3 and WNK4 and provide evidence for the potential role of this PTM in both the established WNK function of osmosensation and a novel function in ferroptosis. While the detailed mechanisms behind these observations remain to be fully elucidated, our results suggest that O-GlcNAc signaling governs the KLHL3/WNK axis at multiple levels and support the broader hypothesis that intracellular glycosylation is a general regulator of the KLHL family and its substrates.

Our MS site-mapping data suggest potential functional consequences for the O-GlcNAcylated residues on KLHL3 and WNK4. We identified T264 as a KLHL3 O-GlcNAc site (Figures 1C and S1A). In a prior study, this residue was also identified as a phosphorylation site, suggesting possible PTM crosstalk (Shibata, S., Arroyo, J.P., et al. 2014). Indeed, crosstalk between phosphorylation and O-GlcNAcylation is a well-documented signaling phenomenon (Wang, Z., Udeshi, N.D., et al. 2010; Hart, G.W., Slawson, C., et al. 2011; Trinidad, J.C., Barkan, D.T., et al. 2012; Wang, S., Huang, X., et al. 2012; Zhong, J., Martinez, M., et al. 2015; Leney, A.C., El Atmioui, D., et al. 2017; Toleman, C.A., Schumacher, M.A., et al. 2018; Schwein, P.A. and Woo, C.M. 2020), motivating future studies to test its potential relevance to KLHL3. T264 is also conserved across multiple species, including zebrafish, mouse, cattle, and non-human primates, suggesting the potential evolutionary importance of this site (Figure S6A). T264 is poorly conserved in other human KLHL proteins, but C265 is conserved in many members (Figure S6B), and the cognate site in KEAP1 (C297) is important for substrate regulation in the oxidative stress response (Dinkova-Kostova, A.T., Holtzclaw, W.D., et al. 2002). Since our results implicate WNK4 in ferroptosis (Figure 5), KLHL3 O-GlcNAcylation could conceivably play a role in oxidative stress responses to ferroptotic or other stimuli by modifying the T264 site adjacent to the conserved C265. Given our serendipitous discovery of WNK O-GlcNAcylation, we focused primarily on WNK4 glycosylation in this work, but dissecting the mechanism and functional consequences of KLHL3 O-GlcNAcylation will be a key goal of future studies.

With respect to WNK4, we observed that S1006 could be either O-GlcNAcylated or phosphorylated (Figures 2F and S1B). In addition, one of the four candidate WNK4 O-GlcNAc sites, S335, is also a known autophosphorylation site (Bazua-Valenti, S., Chavez-Canales, M., et al. 2015; Murillo-de-Ozores, A.R., Rodriguez-Gama, A., et al. 2021). Furthermore, our osmosensation data indicate that phosphorylation of WNK4 S575 may be reduced in the WNK4 6A mutant under hypertonic conditions, relative to WT (Figure S2), suggesting PTM crosstalk in this context. Systematically examining potential O-GlcNAc/O-phosphate crosstalk on WNK4 will be an important goal in the future. We have not yet mapped O-GlcNAc sites on the other three WNK kinases, but sequence alignments indicate that some WNK4 O-GlcNAc sites are serines or threonines at cognate positions in other WNKs (Figure S6C), providing promising candidates to test for O-GlcNAcylation.

Our IP/WB results did not reveal individual O-GlcNAc sites that reduced WNK4 O-GlcNAc levels under basal or TG-induced conditions when mutated (Figure 2G-H). However, both the 2A and 6A mutants failed to increase O-GlcNAcylation in response to TG (Figure 2G-H) and exhibited phenotypes in our osmolarity and ferroptosis experiments (Figures 3 and 5), indicating that the O-GlcNAc sites we identified may together mediate WNK4 responses to these stimuli. Since O-GlcNAc is typically a sub-stoichiometric PTM and MS site-mapping is prone to false-negatives (Udeshi, N.D., Hart, G.W., et al. 2024), it is possible that there are additional WNK4 O-GlcNAc sites awaiting identification.

Functionally, our results revealed an interesting potential connection between osmolarity and WNK4 O-GlcNAcylation in two ways (Figures 3-4). First, hypertonic conditions induced O-GlcNAcylation of WT WNK4 but not of the 6A mutant (Figure 3), suggesting that WNK4’s responses to osmolarity changes may be regulated by its O-GlcNAcylation. Second, and consistent with this hypothesis, the relocalization of WNK4 in response to osmotic changes was altered by chemical inhibition of OGT or OGA, demonstrating that normal O-GlcNAc cycling is required for WNK4 osmosensation (Figure 4). Future studies will be needed to determine whether and which WNK4 O-GlcNAc sites are required for its relocalization and potential phase separation in response to osmolarity changes. Because overexpression of WNKs is known to affect WNK localization and kinase activity in cultured cells and *in vivo* (Lalioti, M.D., Zhang, J., et al. 2006; Tu, S.W., Bugde, A., et al. 2011; Vidal-Petiot, E., Elvira-Matelot, E., et al. 2013), it will likely be necessary to use CRISPR methods to mutate O-GlcNAc sites of interest in the endogenous *WNK4* locus to preserve physiological expression levels and address these questions.

We also discovered a new function of WNK4 as a ferroptosis mediator, consistent with a recent report of a similar role for WNK1 (Xu, X., Sun, Y., et al. 2024). Both our 786-O cell survival assay and OBSC experiments indicate that loss of WNK4 function protects cells from ferroptosis, and WNK4 O-GlcNAcylation may play a role in this process (Figure 5). Interestingly, another recent study suggested that WNK4 knockdown activates the transcription factor STAT3, and STAT3 promotes the expression of GPX4, whose activity protects against ferroptosis (Zhang, W., Gong, M., et al. 2022; Li, M., Shao, X., et al. 2024). While the OBSC system is not amenable to quantitative measurements of STAT3 activation or GPX4 protein expression, future studies will be designed to determine whether WNK4 activity influences ferroptosis through the STAT3/GPX4 pathway and/or other novel interactors to elucidate the molecular mechanisms behind this non-canonical function.

In sum, our study revealed O-GlcNAcylation on KLHL3 and the WNK kinases and identified specific glycosites on KLHL3 and WNK4. These results support our hypothesis that O-GlcNAc regulates the KLHL family and their substrates and expand our knowledge of PTM regulation of WNKs. Our work also highlights the potentially complex crosstalk between WNK O-GlcNAcylation and phosphorylation. O-GlcNAcylation in the KLHL3/WNK axis may be an important regulatory mechanism for multiple processes, including but not limited to osmosensation, ferroptosis, WNK phase separation, and downstream ion channel regulation. The upstream stimuli, pathways, and molecular mechanisms that effect changes in KLHL3/WNK O-GlcNAcylation and the corresponding downstream consequences will be important subjects of future work. We anticipate that our results will inform investigations of KLHL3/WNK pathway regulation and function in different physiological and pathological contexts.

## Materials and Methods

### Cell culture

HEK293T (ATCC CRL-11268) were grown in Dulbecco’s Modified Eagle Medium (DMEM, Sigma, D6429 or Gibco, 11995-065) containing 10% fetal bovine serum (FBS, Sigma, F0926), 100 units/mL penicillin, and 100 µg/mL streptomycin (Pen/Strep, 1% v/v, Gibco, 15140-122). 786-O cells (ATCC CRL-1932) were grown in RPMI 1640 Medium (Gibco, 11875-093) supplemented with 10% FBS and 1% Pen/Strep. Both cell lines were cultured at 37 °C with 5% CO_2_. For osmotic stress experiments, HEK293T cells expressing WNK4-myc-6xHis were cultured for 30 minutes either in hypotonic media: 90 mM NaCl, 2 mM KCl, 1 mM KH_2_PO_4_, 2 mM CaCl_2_, 2 mM MgCl_2_, 10 mM glucose, 10 mM HEPES, pH 7.4; isotonic media: 130 mM NaCl, 2 mM KCl, 1 mM KH_2_PO_4_, 2 mM CaCl_2_, 2 mM MgCl_2_, 10 mM glucose, 20 mM mannitol, 10 mM HEPES, pH 7.4; hypertonic media: 130 mM NaCl, 2 mM KCl, 1 mM KH_2_PO_4_, 2 mM CaCl_2_, 2 mM MgCl_2_, 10 mM glucose, 220 mM mannitol, 10 mM HEPES, pH 7.4.

### Transfections

For all osmotic stress experiments, unless otherwise indicated, cells were transfected with 5 µg of DNA at 50-60% cell density with Lipofectamine 3000 (ThermoFisher, L3000075) for 24 hrs. For other WB experiments and for O-GlcNAc site-mapping, cells were transfected with Mirus TransIT-LT1 (Mirus Bio, MIR 2304) or Lipofectamine 3000 (ThermoFisher, L3000075) in 1:2-1:3 DNA (µg) to transfection reagent (µL) ratio according to manufacturers’ guidelines. DNA plasmids used for transfection and transfection time are indicated in respective figure legends.

### Chemical synthesis

Synthesis of Ac_4_5SGlcNAc (abbreviated as 5SG in text) was done by the Duke Small Molecule Synthesis Facility as described in Gloster et al. (Gloster, T.M., Zandberg, W.F., et al. 2011). All other chemicals were purchased from commercial sources and listed in Table 1.

**Table 1.** Chemical information.

| Chemical name | Vendor and catalog number |
| --- | --- |
| Thiamet-G | Cayman, 13237 |
| OSMI-1 | Cayman, 21894 |
| Erastin | Sigma, 329600 |
| ZT-1a | MedChemExpress, HY-136532 |
| WNK463 | Sigma, SML2809 |
| Fer-1 | Cayman, 17729 |

### IP

Cells were harvested in cold phosphate-buffered saline (PBS) and lysed in cold IP lysis buffer (20 mM Tris-HCl pH 7.4, 1% Triton X-100, 0.1% SDS, 150 mM NaCl, 1 mM EDTA) with protease inhibitor cocktail (PIC, Sigma, P8340, 1:100), 50 µM UDP (Sigma, 94330; OGT inhibitor), 5 µM PUGNAc (Cayman Chemical, 17151; OGA inhibitor), plus 200 µM Na_3_VO_4_ (Sigma, 13721-39-6) for osmotic stress experiments. Lysates were probe-sonicated and cleared by centrifugation at 4 °C. Cleared lysates were quantified by bicinchoninic acid assay (ThermoFisher, 23225). IPs were performed on 2-5 mg of total protein normalized to ∼1-2.5 mg/mL concentration using IP lysis buffer with supplements as described above. Unless otherwise indicated, 2-3 µg of antibody against epitope tags per 1 mg of protein was added to the lysate for rotation overnight at 4 °C. 15-20 µL of settled protein A/G UltraLink resin (ThermoFisher, 53133) was washed three times in the IP lysis buffer and added to each IP either the same time along with antibody for overnight rotation or the next day for rotation at 4 °C for 2 hrs. Beads were washed three times with IP lysis buffer and eluted with SDS-PAGE loading buffer. Either 1X loading buffer supplemented with 5% fresh β-mercaptoethanol (Sigma, M3148) or 2.5X loading buffer was used for elution (5X SDS-PAGE loading buffer: 250 mM Tris pH 6.8, 10% SDS, 30% glycerol, 5% β-mercaptoethanol, 0.02% bromophenol blue). Eluates were heated at 95 °C for 5 min and analyzed by WB.

### WB

For enhanced chemiluminescence (ECL) detection, SDS-PAGE gels were electroblotted onto 100% methanol pre-soaked polyvinylidene difluoride membranes (PVDF, 0.45 µm, ThermoFisher, 88518) in transfer buffer (25 mM Tris, 192 mM glycine, 0.1% SDS, 20% methanol) using a BioRad TransBlot Turbo system. Then, membranes were incubated in blocking buffer (2.5% (w/v) bovine serum albumin (BSA) in Tris-buffered saline with Tween [TBST] [150 mM NaCl, 10 mM Tris-HCl pH 8.0, 0.1% Tween 20]) with agitation at room temperature (RT) for 30 min. Membranes were incubated with primary antibody in blocking buffer overnight at 4 °C. The next day, membranes were washed three times with TBST, each 10 min, and incubated with the appropriate horseradish peroxidase (HRP)-conjugated secondary antibody diluted in blocking buffer at RT for 1 hr. Membranes were again washed three times with TBST, each 10 min. Bands were visualized using ECL (Genesee Scientific, 20-300B) and photographic film (LabScientific, XAR ALF 2025). For quantitative fluorescent WBs, gels were electroblotted onto nitrocellulose membrane (0.45 µm, Bio-Rad, 1620115). Blocking, primary, and washing conditions before secondary antibody incubation were the same as above. Membranes were incubated with appropriate IRDye-conjugated secondary antibody diluted in blocking buffer at RT in the dark for 1 hr. After secondary antibody incubation, membranes were washed three times with TBST, each 10 min, followed by 1-2 TBS washes (TBST but without Tween 20). Bands were visualized using a LI-COR Odyssey CLx Infrared Imaging System. Complete antibody information and dilutions used are provided in Table 2.

**Table 2.** Antibody information.

| Antibody | Vendor and catalog number | Dilutions |
| --- | --- | --- |
| myc (9E10) | BioLegend, 626802 | 1:1000, 1:5000 |
| O-GlcNAc (Multi mAb) | Cell Signaling Technology, 82332 | 1:1000, 1:2000, 1:5000 |
| O-GlcNAc (18B10) | ThermoFisher, MA1-038 | 1:1000 |
| O-GlcNAc (9D1) | ThermoFisher, MA1-039 | 1:1000, 1:2000 |
| O-GlcNAc (RL2)<br>(used for WB and IF) | BioLegend, 677902; Santa Cruz<br>Biotechnology, sc-59624 (discontinued) | 1:1000 (WB), 1:400 (IF) |
| O-GlcNAc (CTD110.6) | Santa Cruz Biotechnology, sc-59623 | 1:200 |
| phospho-WNK4 (Ser575) | ThermoFisher, PA5-114673 | 1:1000 |
| V5 | Cell Signaling Technology, 13202 (IB);<br>ThermoFisher, R960CUS (IP) | 1:10000, 1:1000000 |
| Flag | Sigma, F1804 | 1:1000 |
| α-tubulin | Sigma, T6074 | 1:10000, 1:100000 |
| GAPDH | Santa Cruz Biotechnology, sc-47724 | 1:2000 |
| WNK4 (for WB) | Cell Signaling Technology, 5713 | 1:1000 |
| WNK4 (for IF) | Novus Biologicals, NB600-284 | 1:400 |
| Goat HRP-conjugated anti-mouse IgG secondary<br>antibody | SouthernBiotech, 1030-05 | 1:5000, 1:10000 |
| Goat HRP-conjugated anti-mouse kappa secondary<br>antibody | SouthernBiotech, 1050-05 | 1:5000, 1:10000 |
| Goat HRP-conjugated anti-mouse IgM secondary<br>antibody | SouthernBiotech, 1021-05 | 1:5000 |
| Goat HRP-conjugated anti-rabbit IgG secondary antibody | SouthernBiotech, 4030-05 | 1:5000, 1:10000, 1:20000 |
| Goat HRP-conjugated anti-rabbit light-chain secondary antibody | SouthernBiotech, 4060-05 | 1:10000 |
| Anti-rabbit IgG, HRP-linked Antibody | Cell Signaling Technology, 7074 | 1:1000 |
| Anti-mouse IgG, HRP-linked Antibody | Cell Signaling Technology, 7076 | 1:1000 |
| Goat IRDye 800CW-conjugated anti-mouse IgG (H+L) secondary antibody | Li-Cor, 925-32210 | 1:20000, 1:30000 |
| Goat IRDye 800CW-conjugated anti-rabbit IgG (H+L) secondary antibody | Li-Cor, 925-32211 | 1:20000, 1:30000 |
| Goat Alexa Fluor 488-conjugated-anti-rabbit (H + L) conjugated secondary antibody | ThermoFisher, A-11008 | 1:400 |
| Goat Alexa Fluor 594-conjugated-anti-mouse (H + L) secondary antibody | ThermoFisher, A-11005 | 1:400 |

### Hexosaminidase treatment

Cell pellets were lysed with Tris-Hexo buffer (20 mM Tris-HCl pH 7.4, 5 mM CaCl_2_, 0.5% Triton X-100) supplemented with PIC (Sigma, P8340, 1:100). Lysates were sonicated, centrifuged, and quantified as for IPs (above). Then, lysates were normalized to 1 mg/mL concentration, and 40 µL β-*N*-acetylhexosaminidase_f_ (New England Biolabs, P0721S) was added per 1 mg of protein. Mixtures of protein lysates and hexosaminidase were incubated in a 37 °C water bath for 1 hr. Normalized lysates after the reaction were transferred to new tubes for IP, where antibody and protein A/G UltraLink resin were added, and IPs proceeded as above. Inputs were made with 1X loading buffer using normalized lysates after the reaction and heated at 95 °C for 5 min before WB.

### Sample preparation and tandem IP for O-GlcNAc site-mapping

HEK293T cells were transfected with KLHL3-myc-6xHis or WNK4-myc-6xHis for 24 hrs, then treated with 50 µM Thiamet-G (Cayman Chemical, 13237) plus 4 mM glucosamine (Sigma, G1514) for 24 hrs for KLHL3 site-mapping or treated with DMSO or 50 µM TG for 3 hrs for WNK4 site-mapping. Cells were collected and whole-cell lysates were processed as for Ips (above). Lysates were normalized to 2 mg/mL for 50-100 mg total protein, then incubated with 2 µg myc antibody per 1 mg total protein and 50-100 µL settled protein A/G UltraLink resin overnight at 4 °C. The next day, beads were washed three times with IP lysis buffer without EDTA (20 mM Tris-HCl pH 7.4, 1% Triton X-100, 0.1% SDS, 150 mM NaCl). Washed beads were eluted twice with Ni-NTA wash buffer (300 mM NaCl, 1% Triton X-100, and 10 mM imidazole in 8 M urea/PBS, 500 µL for each elution) by rotation at RT for 15 min per elution. Eluates were then incubated with 40 µL washed, settled HisPur Ni-NTA resin (ThermoFisher, 88223) for 2 hrs RT. After incubation, Ni-NTA resin was washed three times with 1 mL Ni-NTA wash buffer (plus one more no-urea wash for WNK4 samples) and eluted with 250 mM imidazole in 8 M urea/PBS (for WNK4 samples, 250 mM imidazole in PBS) for 3-5 times in 40-80 µL volume by vigorous shaking at RT for 20 min per elution.

### Liquid chromatography-tandem MS analysis

Colloidal blue-stained SDS-PAGE bands (Invitrogen NuPage 4-12% Bis-Tris) were manually excised and subjected to reduction, alkylation, and in-gel tryptic digestion as described (Chen, P.H., Smith, T.J., et al. 2017; Chen, P.H., Hu, J., et al. 2020; Bisnett, B.J., Condon, B.M., et al. 2021; Huynh, D.T., Tsolova, K.N., et al. 2023). Dried samples were subjected to chromatographic separation on a Waters NanoAquity UPLC equipped with a 1.7 μm BEH130 C18 75 μm I.D. × 250 mm reversed-phase column. The mobile phase consisted of (A) 0.1% formic acid (FA) in water and (B) 0.1% FA in acetonitrile. Following a 4 μL injection, peptides were trapped for 3 min on a 5 μm Symmetry C18 180 μm I.D. × 20 mm column at 5 μL/min in 99.9% A. The analytical column was then switched in-line, and a linear elution gradient of 5% B to 40% B was performed over 60 min at 400 nL/min. The analytical column was connected to a fused silica PicoTip emitter (New Objective) with a 10-μm tip orifice and coupled to a Lumos mass spectrometer (ThermoFisher) through an electrospray interface operating in data-dependent acquisition mode. The instrument was set to acquire a precursor MS scan from m/z 350 to 1800 every 3 s. In data-dependent mode, MS/MS scans of the most abundant precursors were collected at r=15,000 (45ms, AGC 5e4) following higher-energy collisional dissociation (HCD) fragmentation at an HCD collision energy of 27%. Within the MS/MS spectra, if any diagnostic O-GlcNAc fragment ions (m/z 204.0867, 138.0545, or 366.1396) were observed, a second MS/MS spectrum at r=30,000 (250ms, 3e5) of the precursor was acquired with electron transfer dissociation (ETD)/HCD fragmentation using charge-dependent ETD reaction times and either 30% (2+ charge state) or 15% (3+-5+ charge state) supplemental collision energy.

For all experiments, a 60 s dynamic exclusion was employed for previously fragmented precursor ions. Raw liquid chromatography–tandem MS (LC-MS/MS) data files were processed in Proteome Discoverer (ThermoFisher) and then submitted to independent Mascot searches (Matrix Science) against a SwissProt database (human taxonomy) containing both forward and reverse entries of each protein (https://www.uniprot.org/proteomes/UP000005640) (20,412 forward entries for KLHL3 MS, 20,348 forward entries for WNK4 MS). Search tolerances were 2 ppm for precursor ions and 0.02 Da for product ions using semi-trypsin specificity with up to two missed cleavages. Both y/b-type HCD and c/z-type ETD fragment ions were allowed for interpreting all spectra. Carbamidomethylation (+57.0214 Da on C) was set as a fixed modification, whereas oxidation (+15.9949 Da on M), phosphorylation (+79.97 Da on S/T) and O-GlcNAc (+203.0794 Da on S/T) were considered dynamic mass modifications. All searched spectra were imported into Scaffold (v4.1, Proteome Software), and scoring thresholds were set to achieve a peptide false discovery rate of 1% using the PeptideProphet algorithm (http://peptideprophet.sourceforge.net/). When satisfactory ETD fragmentation was not obtained upon manual inspection, HCD fragmentation was used to determine O-GlcNAc residue modification using the number of HexNAcs identified in combination with the number of Ser/Thr residues in the peptide.

### Plasmids

Human KLHL3-myc-Flag (DDK) was purchased from OriGene (RC218393), and human WNK4-V5 was purchased from DNASU (HsCD00860697). OGT-myc-6xHis was generated using standard methods as described in (Boyce, M., Carrico, I.S., et al. 2011). KLHL3 and WNK4 were subcloned into pcDNA4-TO-myc-6xHis A using Gibson assembly as described (Huynh, D.T., Tsolova, K.N., et al. 2023). WNK4 S630A, S1006A and 2A were generated from WT WNK4-myc-6xHis using the Q5 Site-Directed Mutagenesis Kit (New England Biolabs, E0552S). WNK4 6A was generated by inserting a gene block containing mutations of S331A, S335A, T543A, and S552A purchased from Integrated DNA Technologies (IDT) into WNK4 2A using Gibson assembly. WNK4 WT and 2A in gWiz backbone were generated from WNK4 WT or 2A in pcDNA4-TO-myc-6xHis A using Gibson assembly. Complete primer sequences and IDT gene block sequence are included in Table 3.

**Table 3.**
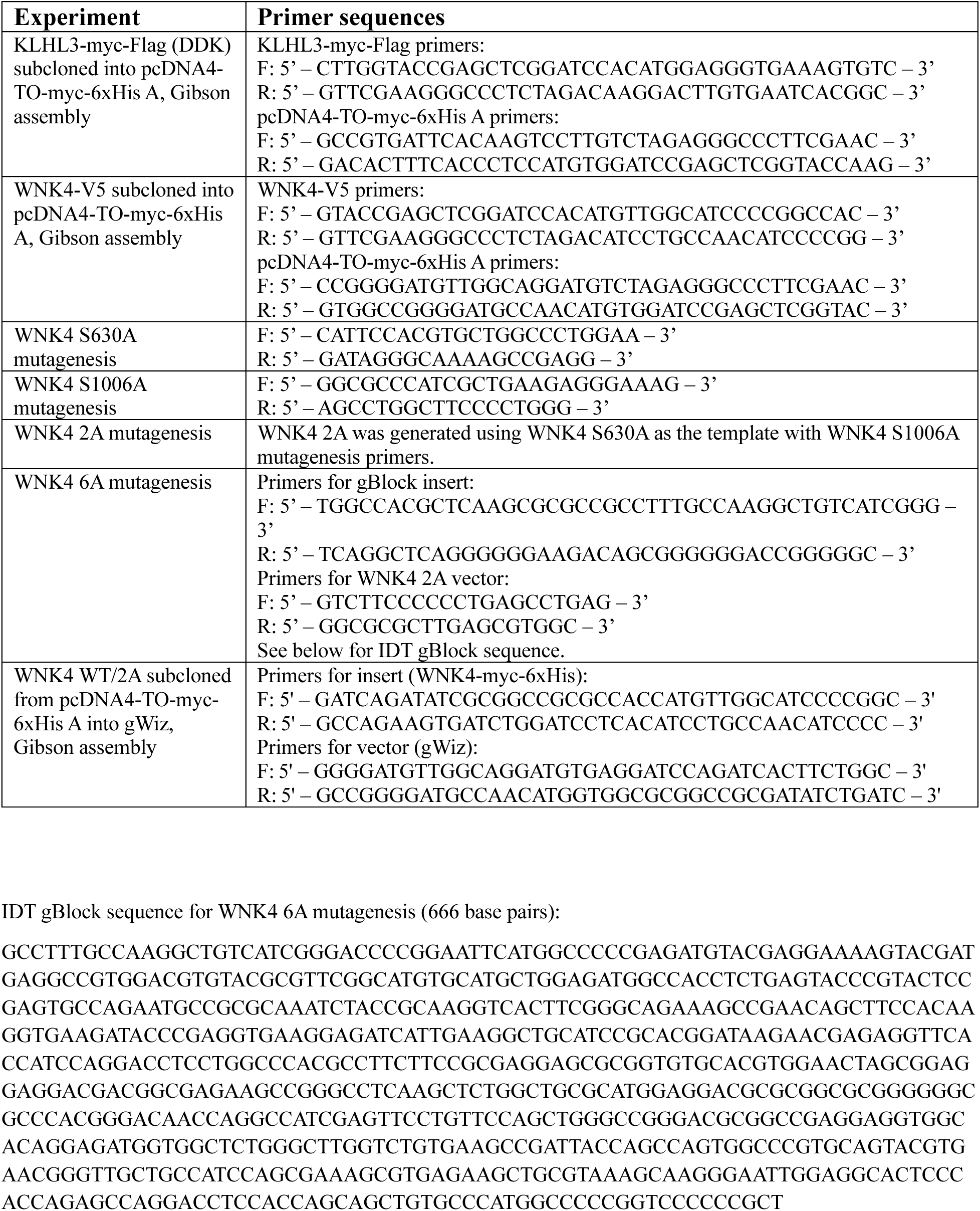
Primer sequences.

### IF

200,000 786-O cells seeded in 12-well plates with an 18 mm coverslip on the bottom were incubated either with DMSO, OSMI-1 (50 µM, 30 min), or TG (50 µM, 30 min). Then, the medium was changed to osmotic buffers (hypotonic or isotonic or hypertonic media – see “Cell culture” section) supplemented either with DMSO, OSMI-1, or TG, and cells were incubated in osmotic buffers for another 30 min. Cells were washed twice with PBS, fixed with 4% paraformaldehyde (MP Biomedicals, 02150146.5, diluted in water) at RT for 15 min, permeabilized with 0.1% Triton X-100/PBS at RT for 10 min, and incubated in blocking buffer (1% BSA/PBS) at RT for 1 hr. Coverslips were incubated with the O-GlcNAc (RL2) and WNK4 antibodies (1:400 in blocking buffer) overnight at 4 °C, washed three times with PBS, and incubated in Alexa Fluor-conjugated secondary antibody (1:400 in blocking buffer) in the dark at RT for 1 hr. Coverslips were washed with PBS twice before mounting in ProLong Diamond anti-fade mounting medium with DAPI (Invitrogen, P36931) onto the microscope slides. Complete antibody information is provided in Table 2.

### Image acquisition on fixed samples

Cells were imaged on an inverted Zeiss 780 single-point scanning confocal microscope equipped with a fully motorized Zeiss Axio Observer microscope base, Marzhauser linearly encoded stage, diode (405 nm), argon ion (488 nm), double solid-state (561 nm), and helium-neon (633 nm) lasers. Images were acquired at RT using a 63x NA/1.4 oil plan apochromatic oil immersion objective lens. Images were acquired sequentially by frame-scanning bidirectionally using the galvanometer-based imaging mode in Zeiss Zen Black acquisition software and processed using Fiji ImageJ. Detection ranges were 480-500 nm, 490-550 nm, and 650-750 nm. Following image acquisition, data were processed using Imaris software (Oxford Instruments) with the “Surface” function to quantify the area for WNK4 immunostaining.

### Cell viability assay

In a 96-well plate, 786-O cells were transfected with a pooled siRNA against WNK4 (Dharmacon) or a non-targeting negative control siRNA using Lipofectamine RNAiMAX Transfection Reagent (ThermoFisher 13778150). After 48 hours, cells were treated with erastin for 24 hours. Cell viability was then evaluated using the CellTiter-Glo Luminescent Cell Viability Assay (Promega G7570) according to manufacturer’s instructions.

Target sequences for WNK4 pooled siRNA:

1. GAUUGCAGCUGCCAUGGUA
2. CAGCUGAGGUGGAGAGUGA
3. GGACAGCUAUGCCUCAGAU
4. CGGGCACGCUCAAGACGUA

### Ferroptotic cell death experiment in OBSCs

Methods for brain slice preparation, biolistic transfection, imaging, and analysis of neuronal health were as described (Braithwaite, S.P., Schmid, R.S., et al. 2010; Dunn, D.E., He, D.N., et al. 2011; Van Kanegan, M.J., Dunn, D.E., et al. 2016). Briefly, hemi-coronal brain slices from postnatal day 8 (P8) rats were prepared and maintained in Neurobasal A medium supplemented with 15% heat-inactivated serum (10% pig serum, 5% rat serum), 10 mM KCl, 10 mM HEPES, 100 U/mL Pen/Strep, 1 mM sodium pyruvate, and 1 mM L-glutamine set in 0.5% reagent-grade agarose. Unless otherwise noted, all brain slices were cultured at 30 °C in humidified incubators under 5% CO_2_. Biolistic transfection of YFP and tau mutant 4R0N was performed by transfecting YFP along with an empty vector or tau4R0N (all constructs in gWiz backbone). Each transfection was performed with 6 mg gold regardless of the total amount of DNA transfected.

For the ZT-1a/WNK463 tau model experiment, OBSCs were sectioned and placed on maintenance medium containing vehicle (DMSO) or the compounds indicated, immediately transfected with 12 µg YFP + 12 µg empty vector or 12 µg YFP + 12µg tau4R0N, and incubated for 72 hrs before visualization. For WNK4 WT/2A experiments, OBSCs were immediately transfected after slices were sectioned and placed in maintenance medium. In addition to 12 µg YFP + 12 µg empty vector (YFP condition) or 12 µg YFP + 12 µg tau4R0N (tau condition), 24 µg empty vector, 24 µg WNK4 WT, or 24 µg WNK4 2A in gWiz backbone was transfected at the same time, and OBSCs were visualized 72 hrs post-transfection. For the ZT-1a/WNK463 stroke model experiment, oxygen-glucose deprivation (OGD) was conducted by suspension of brain slices in glucose-deficient medium (glucose-free, N_2_-bubbled artificial cerebrospinal fluid containing low O_2_ (<0.5%)) at 34 °C for 4.5 min. Control and OGD samples were then plated in maintenance medium containing vehicle (DMSO) or the compounds indicated, transfected with 12 µg YFP, and incubated at 30 °C for 24 hrs before visualization. OBSC visualization was done using Leica MZIIIFL fluorescence stereomicroscope at varied magnifications. Number of biological replicates and statistical tests performed for each experiment are provided in the figure legends.

### Ethical approval

All animal work in this study was performed under the oversight of the Duke University Institutional Animal Care and Use Committee, which reviewed and approved the written protocol (#A233-23-11) and regularly inspects all animal facilities. Rats were acquired through Duke’s Division of Laboratory Animal Resources, which has expert veterinary staff to attend to their daily care and well-being. Rats were housed in a Duke animal care facility accredited by the Association for Assessment and Accreditation of Laboratory Animal Care (AAALAC). All animal procedures, including euthanasia, were carried out according to the general guidelines of the US Animal Welfare Act and AAALAC and complied fully with all relevant ethical standards.

## Statistical analysis

All experimental data presented are representative of multiple independent biological replicates, with replicate numbers indicated in the figure legends. All statistical analyses were performed in Prism 10 with statistical tests and levels of significance explained in figure legends.

## Sequence alignment

For Figure S6A, protein sequences were downloaded from Uniprot (UniProt, C. 2024), and sequence alignment (Clustal Omega 1.2.3 option) and tree construction were completed using Geneious Prime version 2025.0 created by Biomatters (available from https://www.geneious.com). Sequence alignment in Figure S6B-C was performed with Uniprot (UniProt, C. 2024).

## Acknowledgements

We thank members of the Boyce Lab, Dr. Samira Musah, Dr. Titilola Kalejaiye, and David Zhu for insights and help with this project. We thank Dr. Erik Soderblom and the Duke Proteomics and Metabolomics Facility for help with MS experiments and Dr. Benjamin Carlson and the Duke Light Microscopy Core Facility for help with image analysis. This study was supported by NIH grants 5R01GM118847 to M.B., 5R01NS111588 to J.T.C. and M.B., 5U01TR003715 to S.R.F., and R35GM147554 to S.A.M.

## Abbreviations

PTM: post-translational modification
OGT: O-GlcNAc transferase
OGA: O-GlcNAcase
TG: Thiamet-G
5SG: Ac_4_5SGlcNAc
WNK: with-no-lysine
NCC: sodium chloride co-transporter
FHHt: familial hyperkalemic hypertension
WT: wild type
IP: immunoprecipitation
WB: Western blot
IF: immunofluorescence
MS: mass spectrometry
FA: formic acid
HCD: higher-energy collisional dissociation
ETD: electron transfer dissociation
ECL: enhanced chemiluminescence
PVDF: polyvinylidene difluoride
TBST: Tris-buffered saline with Tween
RT: room temperature
HRP: horseradish peroxidase
OBSC: organotypic brain slice cultures
DMEM: Dulbecco’s Modified Eagle Medium
FBS: fetal bovine serum
Pen/Strep: penicillin/streptomycin
LC-MS/MS: liquid chromatography-tandem MS
PIC: protease inhibitor cocktail
PBS: phosphate-buffered saline
BSA: bovine serum albumin
IDT: Integrated DNA Technologies

## Data Availability Statement

KLHL3 and WNK4 O-GlcNAc site-mapping data have been uploaded to the ProteomeXchange Consortium using the PRIDE partner repository with accession number PXD055905.

---

**Figure S1.**
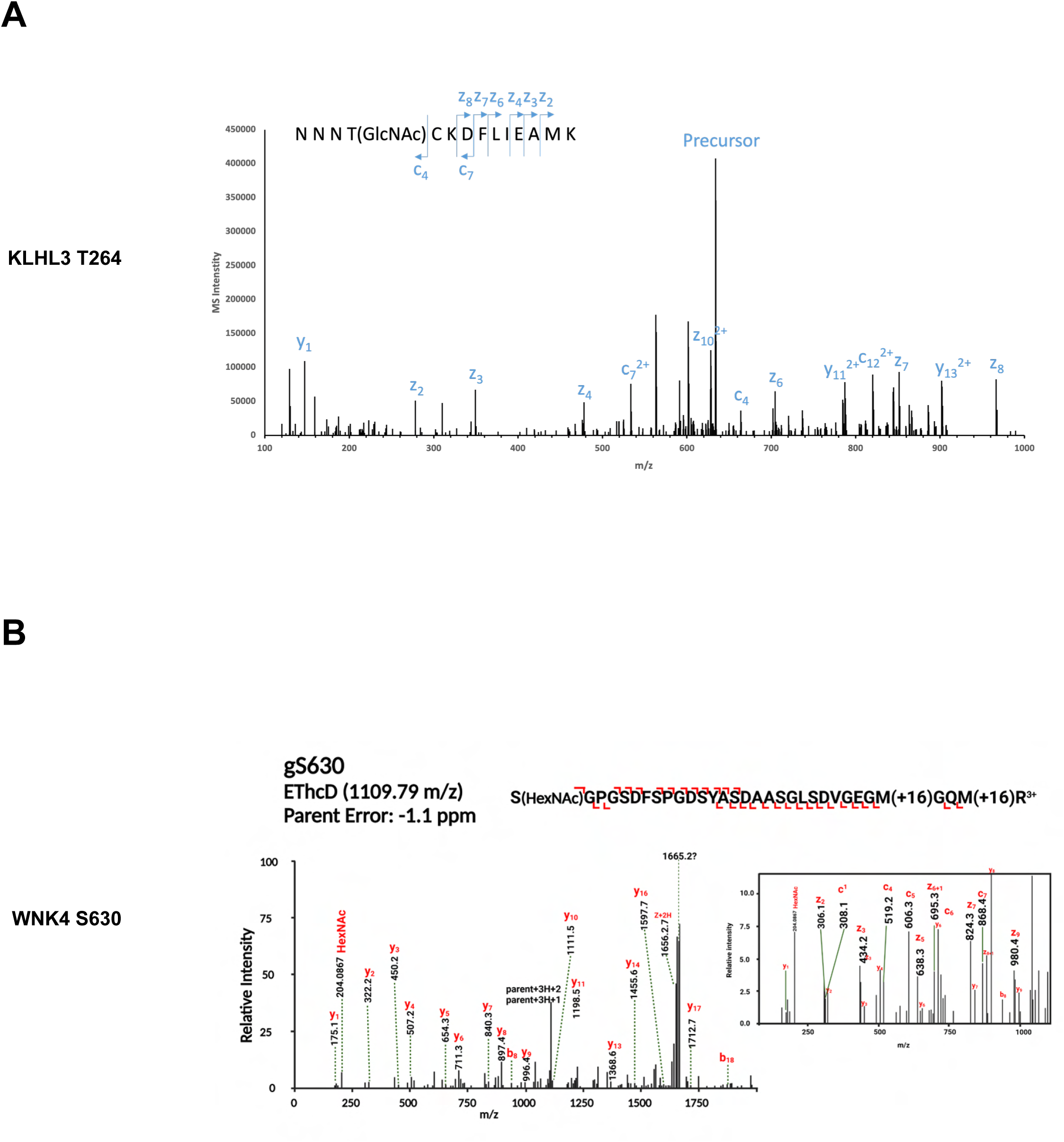
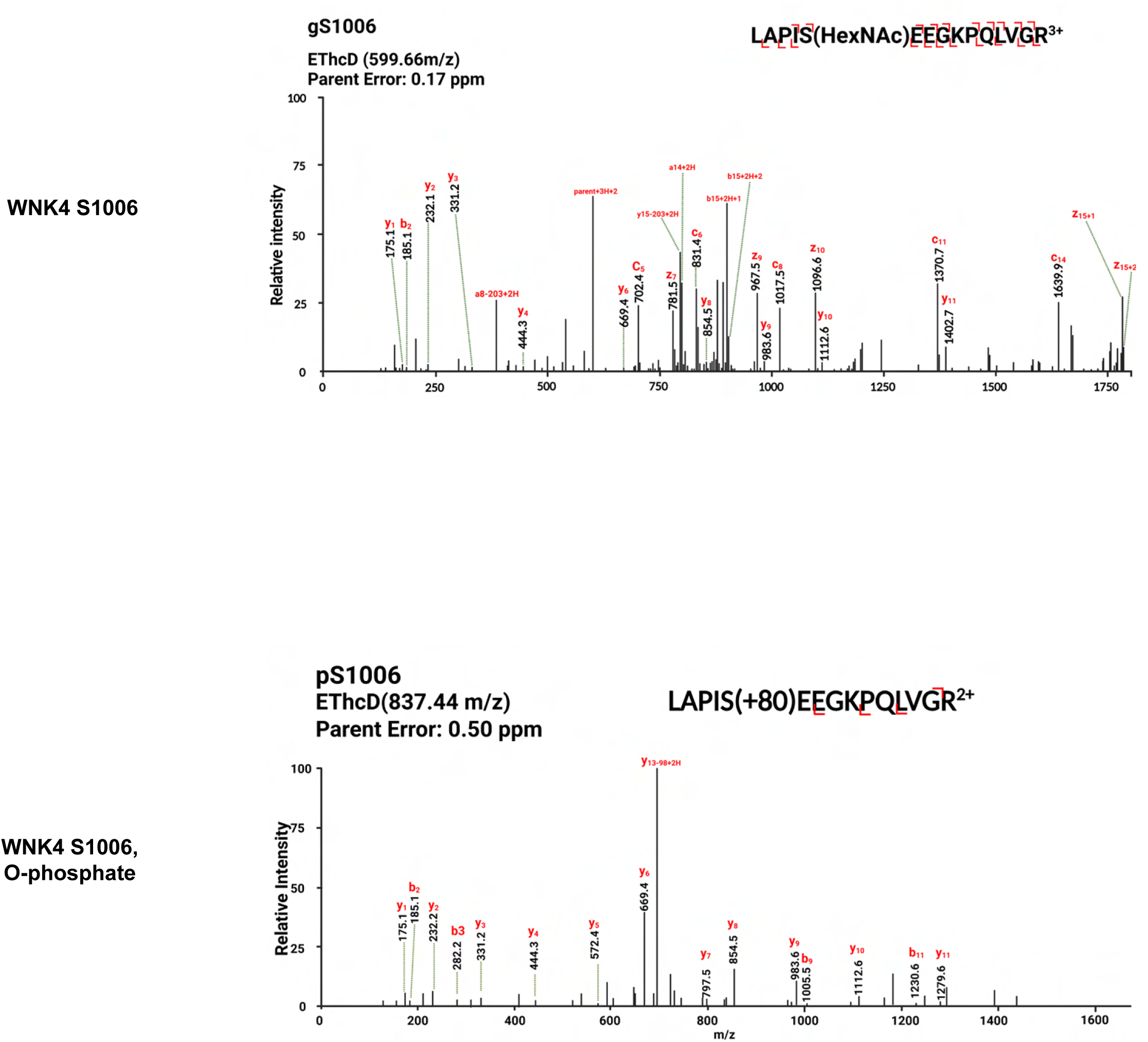
(related to Figures 1C and 2F). (A) Annotated spectrum for MS-identified T264-O-GlcNAc on KLHL3. (B) Annotated spectra for MS-identified S630-O-GlcNAc, S1006-O-GlcNAc. and S1006-O-phosphate on WNK4.

**Figure S2.**
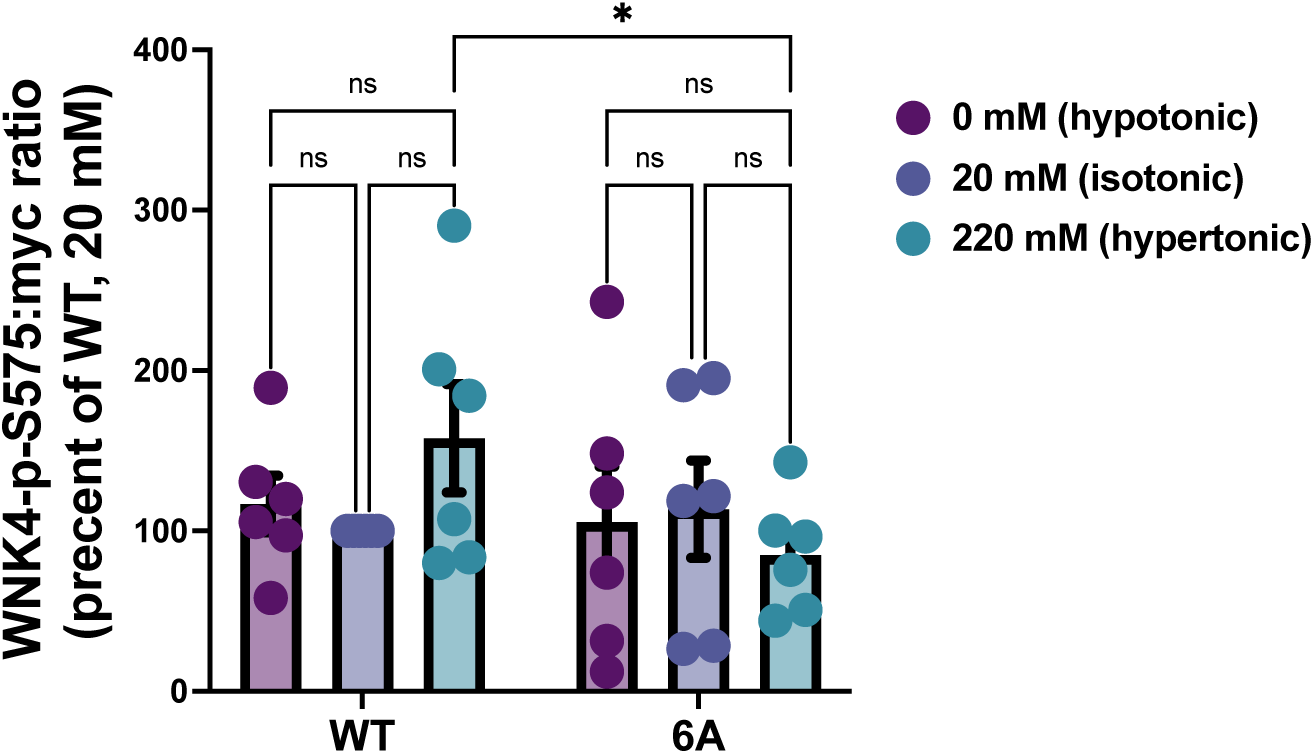
(related to Figure 3A). Fluorescent WBs for total (myc) and phospho-S575 WNK4, depicted in Figure 3A, were quantified and analyzed by two-way ANOVA and Tukey’s multiple comparisons test. All data are displayed as mean ± SEM. n=6. *, *p* ≤ 0.05; ns, not significant.

**Figure S3.**
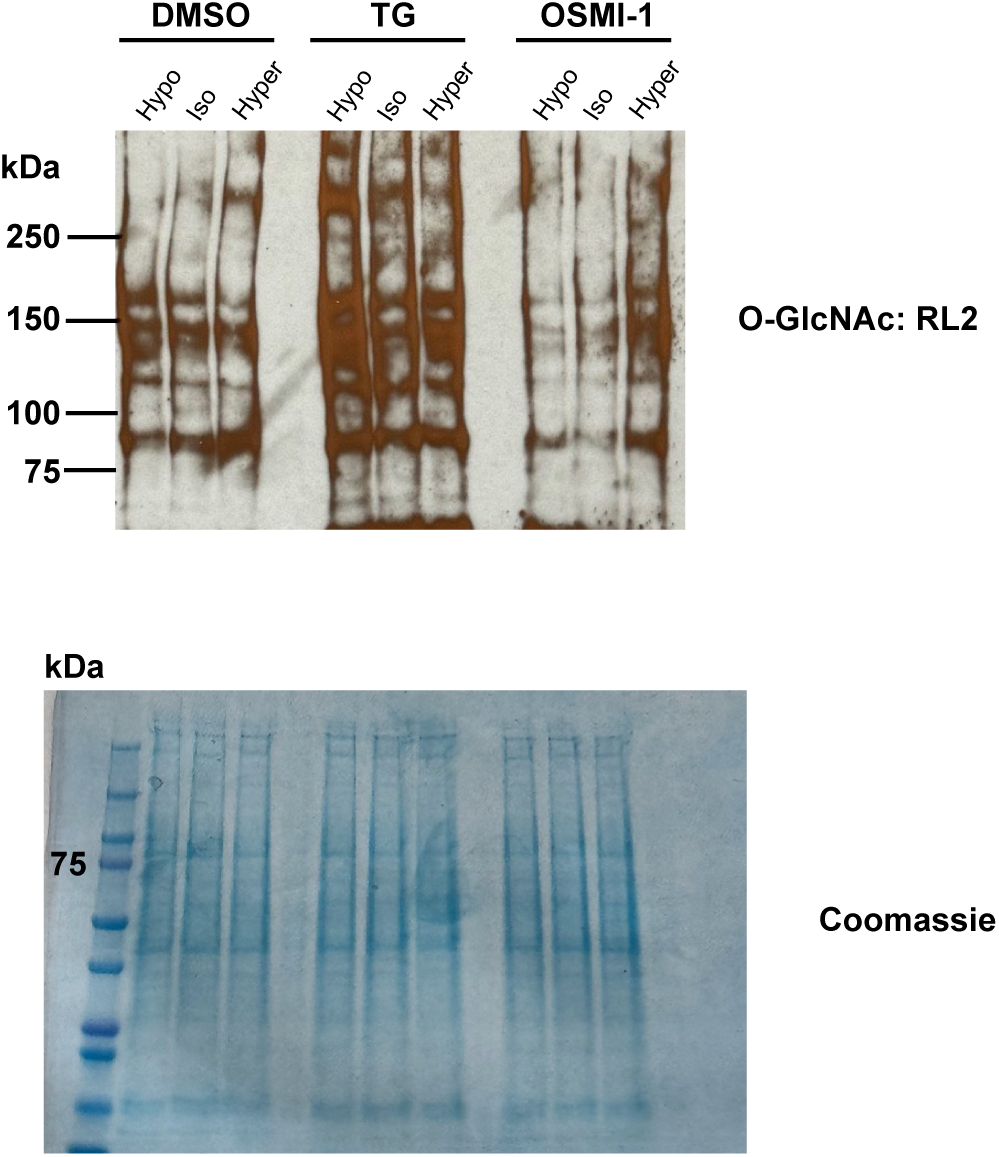
(related to Figure 4). 786-O cells were treated as described in Figure 4, and lysates were analyzed by O-GlcNAc WB and Coomassie stain (loading control).

**Figure S4.**
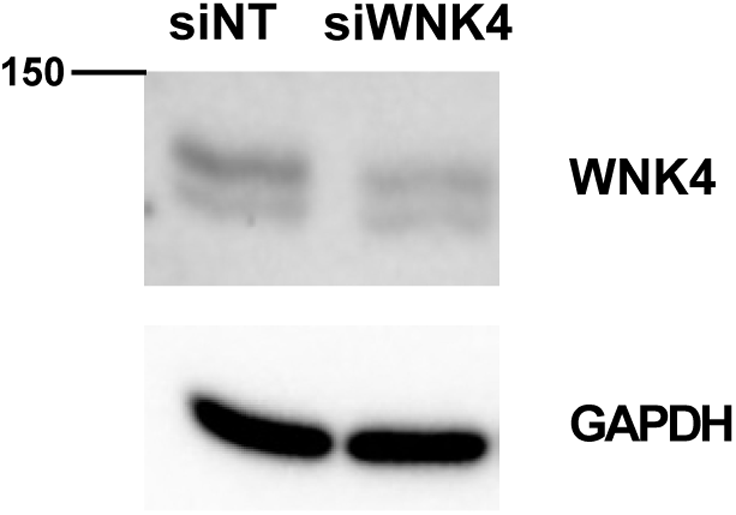
(related to Figure 5A). 786-O cells were treated with control siRNA or anti-WNK4 siRNA, as described in Figure 5A, and lysates were analyzed by WB. GAPDH is a loading control.

**Figure S5.**
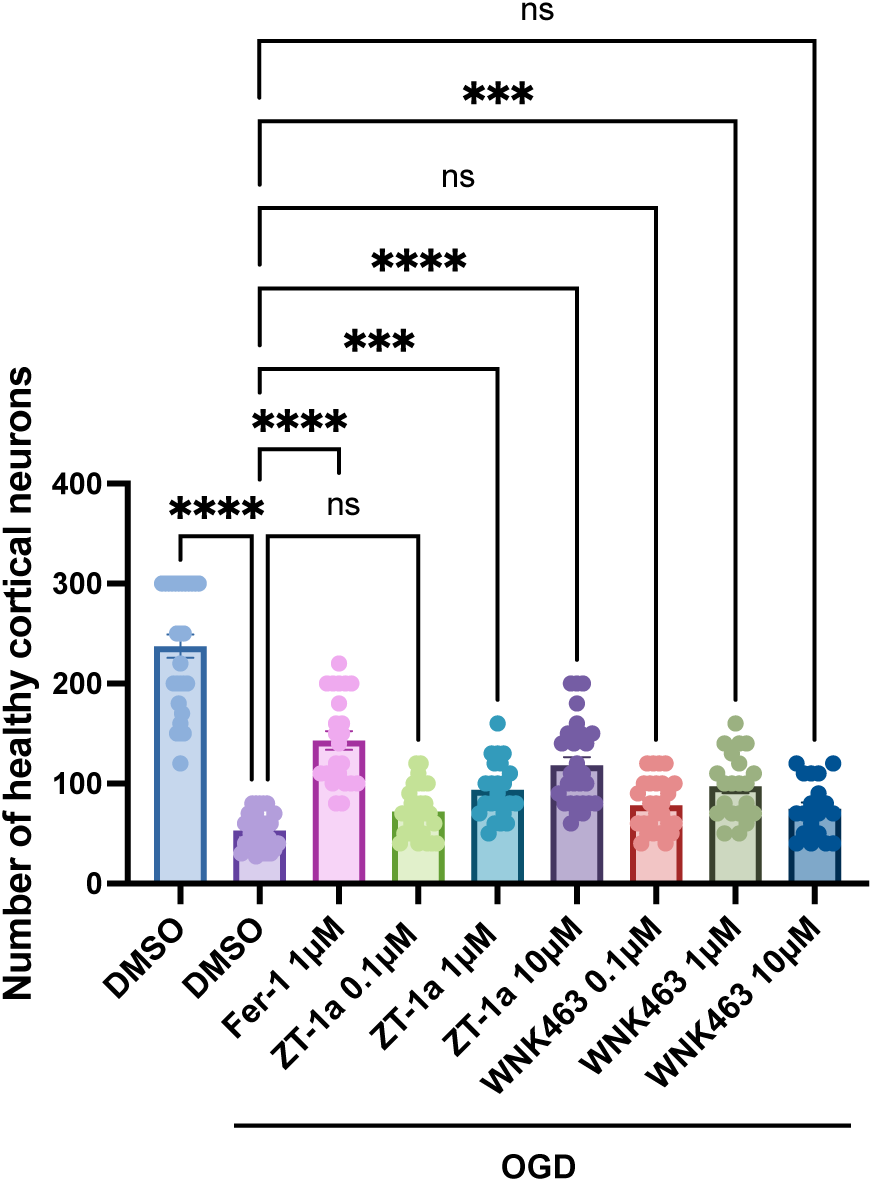
(related to Figure 5B-D). Control or oxygen-glucose deprivation (OGD) samples were treated with DMSO (vehicle), ferrostatin-1 (Fer-1), ZT-1a, or WNK463, as indicated, immediately transfected with YFP, and analyzed by fluorescence microscopy after 24-hour incubation. For each condition, 19-31 brain slices were included for quantitation, the number of healthy cortical neurons on each brain slice was counted and plotted on the graph, and data were analyzed by one-way ANOVA with Dunnett’s multiple comparisons test. All data are displayed as mean ± SEM. n=3. ***, *p* ≤ 0.001; ****, *p* ≤ 0.0001; ns, not significant.

**Figure S6.**
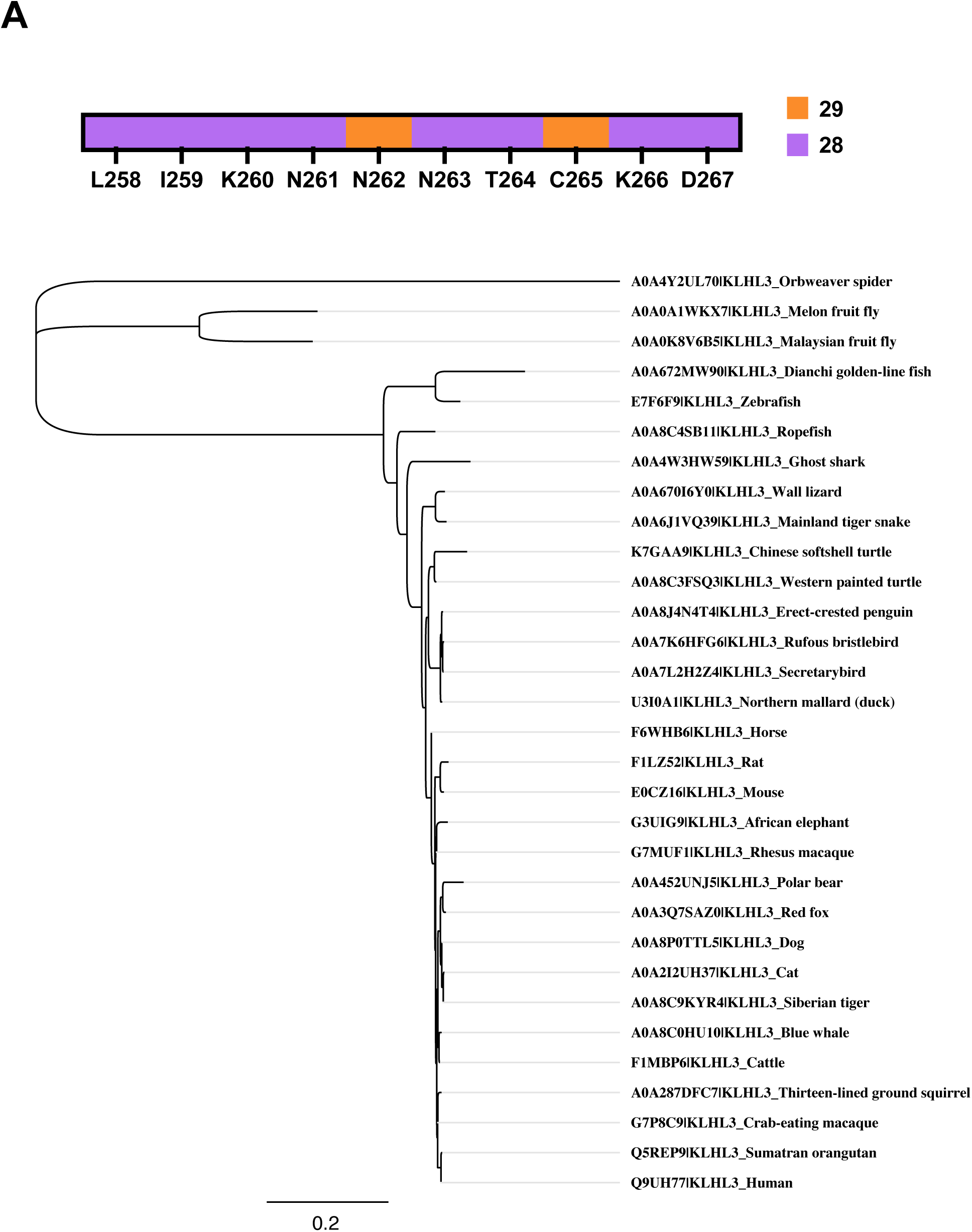
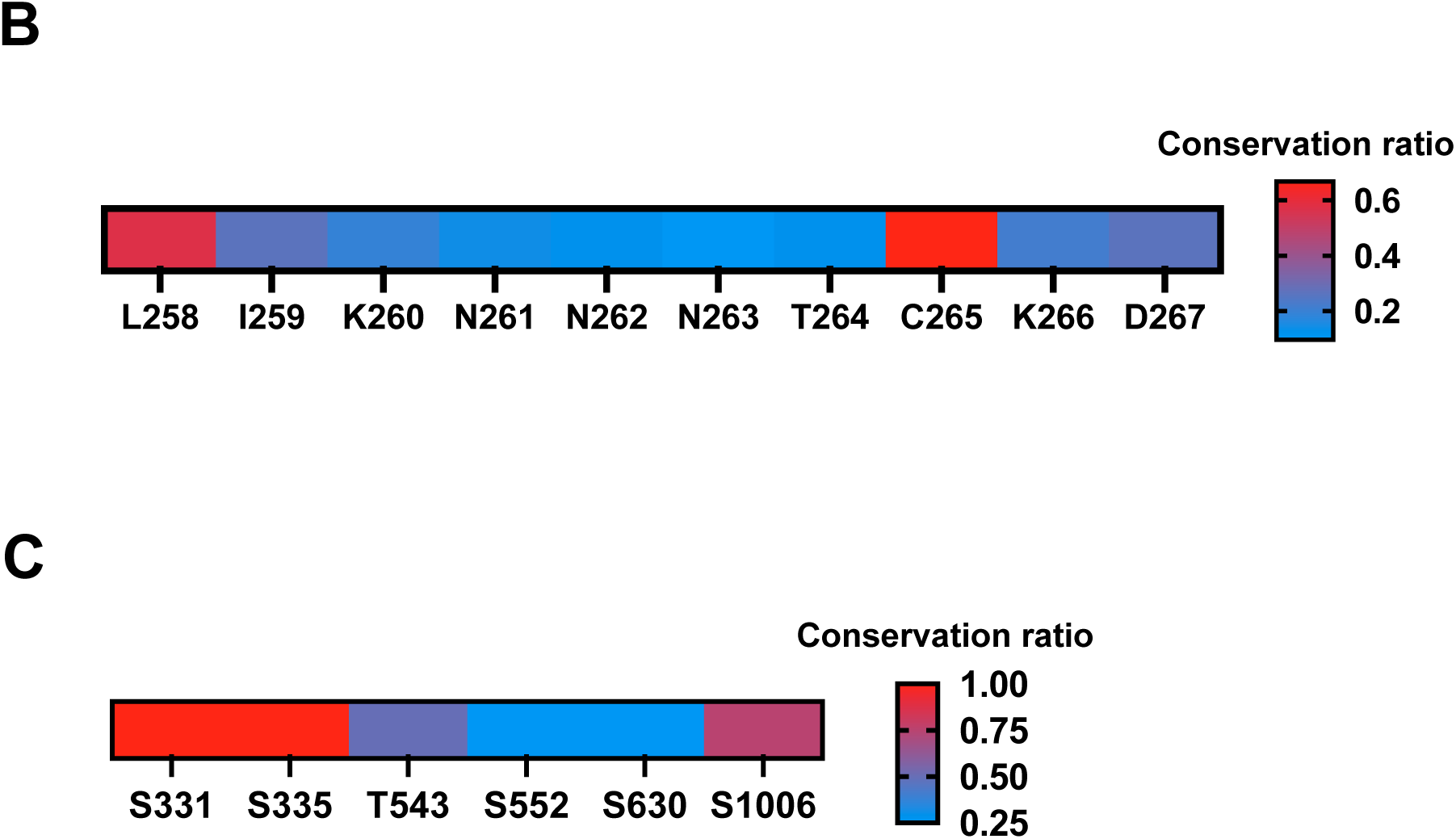
(related to Discussion). (A) KLHL3 sequences from 31 animal species were aligned. Top, heatmap showing the number of species in which human KLHL3 residues are conserved. The human KLHL3 O-GlcNAc site T264 is conserved in 28 of the 31 species selected. Bottom, phylogenetic tree of the 31 species selected. (B) Sequence alignment of 42 human KLHL proteins demonstrates that the KLHL3 O-GlcNAc site T264 is next to a conserved cysteine whose cognate residue in KEAP1 is functionally important in sensing oxidative stress. The number of times that each KLHL3 residue is conserved across all 42 KLHL proteins was divided by 42 to obtain a ratio of conservation, which was then plotted on the heatmap, with red indicating the highest level of conservation and blue indicating the lowest. (C) Sequence alignment of human WNK1-4 suggests potentially conserved O-GlcNAc sites. For each WNK4 O-GlcNAc site, the number of appearances of serine/threonine as cognate residues in WNK1-3 was counted, and a conservation ratio was calculated for each position by dividing the count by 4. Heatmap shows the different frequencies, with red indicating the most conserved and blue indicating the least conserved.

## References

1. Bazua-Valenti S, Chavez-Canales M, Rojas-Vega L, Gonzalez-Rodriguez X, Vazquez N, Rodriguez-Gama A, Argaiz ER, Melo Z, Plata C, Ellison DH, et al. 2015. The Effect of WNK4 on the Na+-Cl-Cotransporter Is Modulated by Intracellular Chloride. J Am Soc Nephrol, 26:1781–1786.

2. Bhuiyan MIH, Young CB, Jahan I, Hasan MN, Fischer S, Meor Azlan NF, Liu M, Chattopadhyay A, Huang H, Kahle KT, et al. 2022. NF-kappaB Signaling-Mediated Activation of WNK-SPAK-NKCC1 Cascade in Worsened Stroke Outcomes of Ang II-Hypertensive Mice. Stroke, 53:1720–1734.

3. Bisnett BJ, Condon BM, Linhart NA, Lamb CH, Huynh DT, Bai J, Smith TJ, Hu J, Georgiou GR, Boyce M. 2021. Evidence for nutrient-dependent regulation of the COPII coat by O-GlcNAcylation. Glycobiology, 31:1102–1120.

4. Boyce M, Carrico IS, Ganguli AS, Yu SH, Hangauer MJ, Hubbard SC, Kohler JJ, Bertozzi CR. 2011. Metabolic cross-talk allows labeling of O-linked beta-N-acetylglucosamine-modified proteins via the N-acetylgalactosamine salvage pathway. Proc Natl Acad Sci U S A, 108:3141–3146.

5. Boyd-Shiwarski CR, Shiwarski DJ, Griffiths SE, Beacham RT, Norrell L, Morrison DE, Wang J, Mann J, Tennant W, Anderson EN, et al. 2022. WNK kinases sense molecular crowding and rescue cell volume via phase separation. Cell, 185:4488–4506 e4420.

6. Boyden LM, Choi M, Choate KA, Nelson-Williams CJ, Farhi A, Toka HR, Tikhonova IR, Bjornson R, Mane SM, Colussi G, et al. 2012. Mutations in kelch-like 3 and cullin 3 cause hypertension and electrolyte abnormalities. Nature, 482:98–102.

7. Braithwaite SP, Schmid RS, He DN, Sung ML, Cho S, Resnick L, Monaghan MM, Hirst WD, Essrich C, Reinhart PH, et al. 2010. Inhibition of c-Jun kinase provides neuroprotection in a model of Alzheimer’s disease. Neurobiol Dis, 39:311–317.

8. Brodaczewska KK, Szczylik C, Fiedorowicz M, Porta C, Czarnecka AM. 2016. Choosing the right cell line for renal cell cancer research. Mol Cancer, 15:83.

9. Chatham JC, Zhang J, Wende AR. 2021. Role of O-Linked N-Acetylglucosamine Protein Modification in Cellular (Patho)Physiology. Physiol Rev, 101:427–493.

10. Chen PH, Chi JT, Boyce M. 2018. Functional crosstalk among oxidative stress and O-GlcNAc signaling pathways. Glycobiology, 28:556–564.

11. Chen PH, Hu J, Wu J, Huynh DT, Smith TJ, Pan S, Bisnett BJ, Smith AB, Lu A, Condon BM, et al. 2020. Gigaxonin glycosylation regulates intermediate filament turnover and may impact giant axonal neuropathy etiology or treatment. JCI Insight, 5.

12. Chen PH, Smith TJ, Wu J, Siesser PF, Bisnett BJ, Khan F, Hogue M, Soderblom E, Tang F, Marks JR, et al. 2017. Glycosylation of KEAP1 links nutrient sensing to redox stress signaling. EMBO J, 36:2233–2250.

13. Chen W, Zebaze LN, Dong J, Chezeau L, Inquimbert P, Hugel S, Niu S, Bihel F, Boutant E, Real E, et al. 2018. WNK1 kinase and its partners Akt, SGK1 and NBC-family Na(+)/HCO3(-) cotransporters are potential therapeutic targets for glioblastoma stem-like cells linked to Bisacodyl signaling. Oncotarget, 9:27197–27219.

14. Chen Y, Zhu G, Liu Y, Wu Q, Zhang X, Bian Z, Zhang Y, Pan Q, Sun F. 2019. O-GlcNAcylated c-Jun antagonizes ferroptosis via inhibiting GSH synthesis in liver cancer. Cell Signal, 63:109384.

15. Chi JT, Lin CC, Lin YT, Chen SY, Setayeshpour Y, Chen Y, Dunn D, Soderblom E, Zhang GF, Filonenko V, et al. unpublished data. https://www.researchsquare.com/article/rs-4522617/v1, last accessed January 24, 2025.

16. Costa RM, Dias MC, Alves JV, Silva JLM, Rodrigues D, Silva JF, Francescato HDC, Ramalho LNZ, Coimbra TM, Tostes RC. 2024. Pharmacological activation of nuclear factor erythroid 2-related factor-2 prevents hyperglycemia-induced renal oxidative damage: Possible involvement of O-GlcNAcylation. Biochem Pharmacol, 220:115982.

17. Cox NJ, Unlu G, Bisnett BJ, Meister TR, Condon BM, Luo PM, Smith TJ, Hanna M, Chhetri A, Soderblom EJ, et al. 2018. Dynamic Glycosylation Governs the Vertebrate COPII Protein Trafficking Pathway. Biochemistry, 57:91–107.

18. Dhanoa BS, Cogliati T, Satish AG, Bruford EA, Friedman JS. 2013. Update on the Kelch-like (KLHL) gene family. Hum Genomics, 7:13.

19. Dinkova-Kostova AT, Holtzclaw WD, Cole RN, Itoh K, Wakabayashi N, Katoh Y, Yamamoto M, Talalay P. 2002. Direct evidence that sulfhydryl groups of Keap1 are the sensors regulating induction of phase 2 enzymes that protect against carcinogens and oxidants. Proc Natl Acad Sci U S A, 99:11908–11913.

20. Dixon SJ, Lemberg KM, Lamprecht MR, Skouta R, Zaitsev EM, Gleason CE, Patel DN, Bauer AJ, Cantley AM, Yang WS, et al. 2012. Ferroptosis: an iron-dependent form of nonapoptotic cell death. Cell, 149:1060–1072.

21. Dixon SJ, Patel DN, Welsch M, Skouta R, Lee ED, Hayano M, Thomas AG, Gleason CE, Tatonetti NP, Slusher BS, et al. 2014. Pharmacological inhibition of cystine-glutamate exchange induces endoplasmic reticulum stress and ferroptosis. Elife, 3:e02523.

22. Dolma S, Lessnick SL, Hahn WC, Stockwell BR. 2003. Identification of genotype-selective antitumor agents using synthetic lethal chemical screening in engineered human tumor cells. Cancer Cell, 3:285–296.

23. Dong DL, Xu ZS, Chevrier MR, Cotter RJ, Cleveland DW, Hart GW. 1993. Glycosylation of mammalian neurofilaments. Localization of multiple O-linked N-acetylglucosamine moieties on neurofilament polypeptides L and M. J Biol Chem, 268:16679–16687.

24. Du L, Wu Y, Fan Z, Li Y, Guo X, Fang Z, Zhang X. 2023. The Role of Ferroptosis in Nervous System Disorders. J Integr Neurosci, 22:19.

25. Dunn DE, He DN, Yang P, Johansen M, Newman RA, Lo DC. 2011. In vitro and in vivo neuroprotective activity of the cardiac glycoside oleandrin from Nerium oleander in brain slice-based stroke models. J Neurochem, 119:805–814.

26. Fei Y, Ding Y. 2024. The role of ferroptosis in neurodegenerative diseases. Front Cell Neurosci, 18:1475934.

27. Ferdaus MZ, McCormick JA. 2018. Mechanisms and controversies in mutant Cul3-mediated familial hyperkalemic hypertension. Am J Physiol Renal Physiol, 314:F915–F920.

28. Fu AB, Xiang SF, He QJ, Ying MD. 2023. Kelch-like proteins in the gastrointestinal tumors. Acta Pharmacol Sin, 44:931–939.

29. Garg A, O’Rourke J, Long C, Doering J, Ravenscroft G, Bezprozvannaya S, Nelson BR, Beetz N, Li L, Chen S, et al. 2014. KLHL40 deficiency destabilizes thin filament proteins and promotes nemaline myopathy. J Clin Invest, 124:3529–3539.

30. Genschik P, Sumara I, Lechner E. 2013. The emerging family of CULLIN3-RING ubiquitin ligases (CRL3s): cellular functions and disease implications. EMBO J, 32:2307–2320.

31. Gloster TM, Zandberg WF, Heinonen JE, Shen DL, Deng L, Vocadlo DJ. 2011. Hijacking a biosynthetic pathway yields a glycosyltransferase inhibitor within cells. Nat Chem Biol, 7:174–181.

32. Goldsmith EJ, Rodan AR. 2023. Intracellular Ion Control of WNK Signaling. Annu Rev Physiol, 85:383–406.

33. Hadchouel J, Ellison DH, Gamba G. 2016. Regulation of Renal Electrolyte Transport by WNK and SPAK-OSR1 Kinases. Annu Rev Physiol, 78:367–389.

34. Hart GW. 2014. Three Decades of Research on O-GlcNAcylation – A Major Nutrient Sensor That Regulates Signaling, Transcription and Cellular Metabolism. Front Endocrinol (Lausanne), 5:183.

35. Hart GW, Slawson C, Ramirez-Correa G, Lagerlof O. 2011. Cross talk between O-GlcNAcylation and phosphorylation: roles in signaling, transcription, and chronic disease. Annu Rev Biochem, 80:825–858.

36. Hoffstrom BG, Kaplan A, Letso R, Schmid RS, Turmel GJ, Lo DC, Stockwell BR. 2010. Inhibitors of protein disulfide isomerase suppress apoptosis induced by misfolded proteins. Nat Chem Biol, 6:900–906.

37. Hu CW, Wang K, Jiang J. 2024. The non-catalytic domains of O-GlcNAc cycling enzymes present new opportunities for function-specific control. Curr Opin Chem Biol, 81:102476.

38. Huynh DT, Tsolova KN, Watson AJ, Khal SK, Green JR, Li D, Hu J, Soderblom EJ, Chi JT, Evans CS, et al. 2023. O-GlcNAcylation regulates neurofilament-light assembly and function and is perturbed by Charcot-Marie-Tooth disease mutations. Nat Commun, 14:6558.

39. Ishibashi K, Koguchi T, Matsuoka K, Onagi A, Tanji R, Takinami-Honda R, Hoshi S, Onoda M, Kurimura Y, Hata J, et al. 2018. Interleukin-6 induces drug resistance in renal cell carcinoma. Fukushima J Med Sci, 64:103–110.

40. Ishizawa K, Wang Q, Li J, Yamazaki O, Tamura Y, Fujigaki Y, Uchida S, Lifton RP, Shibata S. 2019. Calcineurin dephosphorylates Kelch-like 3, reversing phosphorylation by angiotensin II and regulating renal electrolyte handling. Proc Natl Acad Sci U S A, 116:3155–3160.

41. Izadifar A, Courchet J, Virga DM, Verreet T, Hamilton S, Ayaz D, Misbaer A, Vandenbogaerde S, Monteiro L, Petrovic M, et al. 2021. Axon morphogenesis and maintenance require an evolutionary conserved safeguard function of Wnk kinases antagonizing Sarm and Axed. Neuron, 109:2864–2883 e2868.

42. Jakaria M, Belaidi AA, Bush AI, Ayton S. 2021. Ferroptosis as a mechanism of neurodegeneration in Alzheimer’s disease. J Neurochem, 159:804–825.

43. Jiang X, Stockwell BR, Conrad M. 2021. Ferroptosis: mechanisms, biology and role in disease. Nat Rev Mol Cell Biol, 22:266–282.

44. Lalioti MD, Zhang J, Volkman HM, Kahle KT, Hoffmann KE, Toka HR, Nelson-Williams C, Ellison DH, Flavell R, Booth CJ, et al. 2006. Wnk4 controls blood pressure and potassium homeostasis via regulation of mass and activity of the distal convoluted tubule. Nat Genet, 38:1124–1132.

45. Le Minh G, Esquea EM, Young RG, Huang J, Reginato MJ. 2023. On a sugar high: Role of O-GlcNAcylation in cancer. J Biol Chem, 299:105344.

46. Leney AC, El Atmioui D, Wu W, Ovaa H, Heck AJR. 2017. Elucidating crosstalk mechanisms between phosphorylation and O-GlcNAcylation. Proc Natl Acad Sci U S A, 114:E7255–E7261.

47. Li M, Shao X, Ning Q, Sun R, Li R, Liu Y, Yuan Y, Zhang Y. 2024. Downregulation of WNK4 expression facilitates the proliferation of gastric cancer cells via activation of the STAT3 signaling pathway. Neoplasma, 71:209–218.

48. Li Y, Xiao D, Wang X. 2022. The emerging roles of ferroptosis in cells of the central nervous system. Front Neurosci, 16:1032140.

49. Liang D, Minikes AM, Jiang X. 2022. Ferroptosis at the intersection of lipid metabolism and cellular signaling. Mol Cell, 82:2215–2227.

50. Louis-Dit-Picard H, Barc J, Trujillano D, Miserey-Lenkei S, Bouatia-Naji N, Pylypenko O, Beaurain G, Bonnefond A, Sand O, Simian C, et al. 2012. KLHL3 mutations cause familial hyperkalemic hypertension by impairing ion transport in the distal nephron. Nat Genet, 44:456–460, S451-453.

51. Lu B, Chen XB, Ying MD, He QJ, Cao J, Yang B. 2017. The Role of Ferroptosis in Cancer Development and Treatment Response. Front Pharmacol, 8:992.

52. Maeoka Y, Cornelius RJ, McCormick JA. 2023. Cullin 3 and Blood Pressure Regulation: Insights From Familial Hyperkalemic Hypertension. Hypertension, 80:912–923.

53. Magnelli P, Bielik A, Guthrie E. 2012. Identification and characterization of protein glycosylation using specific endo– and exoglycosidases. Methods Mol Biol, 801:189–211.

54. Maruyama J, Kobayashi Y, Umeda T, Vandewalle A, Takeda K, Ichijo H, Naguro I. 2016. Osmotic stress induces the phosphorylation of WNK4 Ser575 via the p38MAPK-MK pathway. Sci Rep, 6:18710.

55. Mayfield JM, Hitefield NL, Czajewski I, Vanhye L, Holden L, Morava E, van Aalten DMF, Wells L. 2024. O-GlcNAc transferase congenital disorder of glycosylation (OGT-CDG): Potential mechanistic targets revealed by evaluating the OGT interactome. J Biol Chem, 300:107599.

56. Murillo-de-Ozores AR, Chavez-Canales M, de Los Heros P, Gamba G, Castaneda-Bueno M. 2020. Physiological Processes Modulated by the Chloride-Sensitive WNK-SPAK/OSR1 Kinase Signaling Pathway and the Cation-Coupled Chloride Cotransporters. Front Physiol, 11:585907.

57. Murillo-de-Ozores AR, Rodriguez-Gama A, Carbajal-Contreras H, Gamba G, Castaneda-Bueno M. 2021. WNK4 kinase: from structure to physiology. Am J Physiol Renal Physiol, 320:F378–F403.

58. Myers SA, Daou S, Affar el B, Burlingame A. 2013. Electron transfer dissociation (ETD): the mass spectrometric breakthrough essential for O-GlcNAc protein site assignments-a study of the O-GlcNAcylated protein host cell factor C1. Proteomics, 13:982–991.

59. Myers SA, Peddada S, Chatterjee N, Friedrich T, Tomoda K, Krings G, Thomas S, Maynard J, Broeker M, Thomson M, et al. 2016. SOX2 O-GlcNAcylation alters its protein-protein interactions and genomic occupancy to modulate gene expression in pluripotent cells. Elife, 5:e10647.

60. Ohta A, Schumacher FR, Mehellou Y, Johnson C, Knebel A, Macartney TJ, Wood NT, Alessi DR, Kurz T. 2013. The CUL3-KLHL3 E3 ligase complex mutated in Gordon’s hypertension syndrome interacts with and ubiquitylates WNK isoforms: disease-causing mutations in KLHL3 and WNK4 disrupt interaction. Biochem J, 451:111–122.

61. Ong Q, Han W, Yang X. 2018. O-GlcNAc as an Integrator of Signaling Pathways. Front Endocrinol (Lausanne), 9:599.

62. Ortiz-Meoz RF, Jiang J, Lazarus MB, Orman M, Janetzko J, Fan C, Duveau DY, Tan ZW, Thomas CJ, Walker S. 2015. A small molecule that inhibits OGT activity in cells. ACS Chem Biol, 10:1392–1397.

63. Park HW, Kim JY, Choi SK, Lee YH, Zeng W, Kim KH, Muallem S, Lee MG. 2011. Serine-threonine kinase with-no-lysine 4 (WNK4) controls blood pressure via transient receptor potential canonical 3 (TRPC3) in the vasculature. Proc Natl Acad Sci U S A, 108:10750–10755.

64. Pratt MR, Vocadlo DJ. 2023. Understanding and exploiting the roles of O-GlcNAc in neurodegenerative diseases. J Biol Chem, 299:105411.

65. Ramirez-Martinez A, Cenik BK, Bezprozvannaya S, Chen B, Bassel-Duby R, Liu N, Olson EN. 2017. KLHL41 stabilizes skeletal muscle sarcomeres by nonproteolytic ubiquitination. Elife, 6.

66. Rao X, Duan X, Mao W, Li X, Li Z, Li Q, Zheng Z, Xu H, Chen M, Wang PG, et al. 2015. O-GlcNAcylation of G6PD promotes the pentose phosphate pathway and tumor growth. Nat Commun, 6:8468.

67. Reinhart PH, Kaltenbach LS, Essrich C, Dunn DE, Eudailey JA, DeMarco CT, Turmel GJ, Whaley JC, Wood A, Cho S, et al. 2011. Identification of anti-inflammatory targets for Huntington’s disease using a brain slice-based screening assay. Neurobiol Dis, 43:248–256.

68. Sanchez-Fdez A, Matilla-Almazan S, Montero JC, Del Carmen S, Abad M, Garcia-Alonso S, Bhattacharya S, Calar K, de la Puente P, Ocana A, et al. 2023. The WNK1-ERK5 route plays a pathophysiological role in ovarian cancer and limits therapeutic efficacy of trametinib. Clin Transl Med, 13:e1217.

69. Satcher RL, Pan T, Cheng CJ, Lee YC, Lin SC, Yu G, Li X, Hoang AG, Tamboli P, Jonasch E, et al. 2014. Cadherin-11 in renal cell carcinoma bone metastasis. PLoS One, 9:e89880.

70. Schwein PA, Woo CM. 2020. The O-GlcNAc Modification on Kinases. ACS Chem Biol, 15:602–617.

71. Sevilla-Montero J, Bienes-Martinez R, Labrousse-Arias D, Fuertes-Yebra E, Ordonez A, Calzada MJ. 2020. pVHL-mediated regulation of the anti-angiogenic protein thrombospondin-1 decreases migration of Clear Cell Renal Carcinoma Cell Lines. Sci Rep, 10:1175.

72. Shaharabany M, Holtzman EJ, Mayan H, Hirschberg K, Seger R, Farfel Z. 2008. Distinct pathways for the involvement of WNK4 in the signaling of hypertonicity and EGF. FEBS J, 275:1631–1642.

73. Shekarabi M, Zhang J, Khanna AR, Ellison DH, Delpire E, Kahle KT. 2017. WNK Kinase Signaling in Ion Homeostasis and Human Disease. Cell Metab, 25:285–299.

74. Shibata S, Arroyo JP, Castaneda-Bueno M, Puthumana J, Zhang J, Uchida S, Stone KL, Lam TT, Lifton RP. 2014. Angiotensin II signaling via protein kinase C phosphorylates Kelch-like 3, preventing WNK4 degradation. Proc Natl Acad Sci U S A, 111:15556–15561.

75. Shibata S, Zhang J, Puthumana J, Stone KL, Lifton RP. 2013. Kelch-like 3 and Cullin 3 regulate electrolyte homeostasis via ubiquitination and degradation of WNK4. Proc Natl Acad Sci U S A, 110:7838–7843.

76. Shimizu M, Shibuya H. 2022. WNK1/HSN2 mediates neurite outgrowth and differentiation via a OSR1/GSK3beta-LHX8 pathway. Sci Rep, 12:15858.

77. Skouta R, Dixon SJ, Wang J, Dunn DE, Orman M, Shimada K, Rosenberg PA, Lo DC, Weinberg JM, Linkermann A, et al. 2014. Ferrostatins inhibit oxidative lipid damage and cell death in diverse disease models. J Am Chem Soc, 136:4551–4556.

78. Slawson C, Lakshmanan T, Knapp S, Hart GW. 2008. A mitotic GlcNAcylation/phosphorylation signaling complex alters the posttranslational state of the cytoskeletal protein vimentin. Mol Biol Cell, 19:4130–4140.

79. Sodi VL, Bacigalupa ZA, Ferrer CM, Lee JV, Gocal WA, Mukhopadhyay D, Wellen KE, Ivan M, Reginato MJ. 2018. Nutrient sensor O-GlcNAc transferase controls cancer lipid metabolism via SREBP-1 regulation. Oncogene, 37:924–934.

80. Stockwell BR, Friedmann Angeli JP, Bayir H, Bush AI, Conrad M, Dixon SJ, Fulda S, Gascon S, Hatzios SK, Kagan VE, et al. 2017. Ferroptosis: A Regulated Cell Death Nexus Linking Metabolism, Redox Biology, and Disease. Cell, 171:273–285.

81. Tang BL. 2016. (WNK)ing at death: With-no-lysine (Wnk) kinases in neuropathies and neuronal survival. Brain Res Bull, 125:92–98.

82. Tarbet HJ, Dolat L, Smith TJ, Condon BM, O’Brien ET, 3rd, Valdivia RH, Boyce M. 2018. Site-specific glycosylation regulates the form and function of the intermediate filament cytoskeleton. Elife, 7.

83. Toleman CA, Schumacher MA, Yu SH, Zeng W, Cox NJ, Smith TJ, Soderblom EJ, Wands AM, Kohler JJ, Boyce M. 2018. Structural basis of O-GlcNAc recognition by mammalian 14-3-3 proteins. Proc Natl Acad Sci U S A, 115:5956–5961.

84. Trinidad JC, Barkan DT, Gulledge BF, Thalhammer A, Sali A, Schoepfer R, Burlingame AL. 2012. Global identification and characterization of both O-GlcNAcylation and phosphorylation at the murine synapse. Mol Cell Proteomics, 11:215–229.

85. Tu SW, Bugde A, Luby-Phelps K, Cobb MH. 2011. WNK1 is required for mitosis and abscission. Proc Natl Acad Sci U S A, 108:1385–1390.

86. Uchida S, Mori T, Susa K, Sohara E. 2022. NCC regulation by WNK signal cascade. Front Physiol, 13:1081261.

87. Udeshi ND, Hart GW, Slawson C. 2024. From Fringe to the Mainstream: How ETD MS Brought O-GlcNAc to the Masses. Mol Cell Proteomics, 23:100859.

88. Umapathi P, Aggarwal A, Zahra F, Narayanan B, Zachara NE. 2024. The multifaceted role of intracellular glycosylation in cytoprotection and heart disease. J Biol Chem, 300:107296.

89. UniProt C. 2024. UniProt: the Universal Protein Knowledgebase in 2025. Nucleic Acids Res.

90. Van Kanegan MJ, Dunn DE, Kaltenbach LS, Shah B, He DN, McCoy DD, Yang P, Peng J, Shen L, Du L, et al. 2016. Dual activities of the anti-cancer drug candidate PBI-05204 provide neuroprotection in brain slice models for neurodegenerative diseases and stroke. Sci Rep, 6:25626.

91. Venkatesh D, O’Brien NA, Zandkarimi F, Tong DR, Stokes ME, Dunn DE, Kengmana ES, Aron AT, Klein AM, Csuka JM, et al. 2020. MDM2 and MDMX promote ferroptosis by PPARalpha-mediated lipid remodeling. Genes Dev, 34:526–543.

92. Vidal-Petiot E, Elvira-Matelot E, Mutig K, Soukaseum C, Baudrie V, Wu S, Cheval L, Huc E, Cambillau M, Bachmann S, et al. 2013. WNK1-related Familial Hyperkalemic Hypertension results from an increased expression of L-WNK1 specifically in the distal nephron. Proc Natl Acad Sci U S A, 110:14366–14371.

93. Wakabayashi M, Mori T, Isobe K, Sohara E, Susa K, Araki Y, Chiga M, Kikuchi E, Nomura N, Mori Y, et al. 2013. Impaired KLHL3-mediated ubiquitination of WNK4 causes human hypertension. Cell Rep, 3:858–868.

94. Wang JKT, Portbury S, Thomas MB, Barney S, Ricca DJ, Morris DL, Warner DS, Lo DC. 2006. Cardiac glycosides provide neuroprotection against ischemic stroke: discovery by a brain slice-based compound screening platform. Proc Natl Acad Sci U S A, 103:10461–10466.

95. Wang S, Huang X, Sun D, Xin X, Pan Q, Peng S, Liang Z, Luo C, Yang Y, Jiang H, et al. 2012. Extensive crosstalk between O-GlcNAcylation and phosphorylation regulates Akt signaling. PLoS One, 7:e37427.

96. Wang Z, Udeshi ND, Slawson C, Compton PD, Sakabe K, Cheung WD, Shabanowitz J, Hunt DF, Hart GW. 2010. Extensive crosstalk between O-GlcNAcylation and phosphorylation regulates cytokinesis. Sci Signal, 3:ra2.

97. Wang ZL, Yuan L, Li W, Li JY. 2022. Ferroptosis in Parkinson’s disease: glia-neuron crosstalk. Trends Mol Med, 28:258–269.

98. Wilson FH, Disse-Nicodeme S, Choate KA, Ishikawa K, Nelson-Williams C, Desitter I, Gunel M, Milford DV, Lipkin GW, Achard JM, et al. 2001. Human hypertension caused by mutations in WNK kinases. Science, 293:1107–1112.

99. Xu X, Sun Y, Zhu X, Ma S, Wei J, He C, Chen J, Pan X. 2024. Lnc_011797 promotes ferroptosis and aggravates white matter lesions. Neural Regen Res.

100. Xu ZQ, Zhang L, Gao BS, Wan YG, Zhang XH, Chen B, Wang YT, Sun N, Fu YW. 2015. EZH2 promotes tumor progression by increasing VEGF expression in clear cell renal cell carcinoma. Clin Transl Oncol, 17:41–49.

101. Yamada K, Park HM, Rigel DF, DiPetrillo K, Whalen EJ, Anisowicz A, Beil M, Berstler J, Brocklehurst CE, Burdick DA, et al. 2016. Small-molecule WNK inhibition regulates cardiovascular and renal function. Nat Chem Biol, 12:896–898.

102. Yang L, Tang H, Wang J, Xu D, Xuan R, Xie S, Xu P, Li X. 2025. O-GlcNAcylation attenuates ischemia-reperfusion-induced pulmonary epithelial cell ferroptosis via the Nrf2/G6PDH pathway. BMC Biol, 23:32.

103. Yang T, Zhao K, Shu H, Chen X, Cheng J, Li S, Zhao Z, Kuang Y, Yu S. 2017. The Nogo receptor inhibits proliferation, migration and axonal extension by transcriptionally regulating WNK1 in PC12 cells. Neuroreport, 28:533–539.

104. Yang X, Qian K. 2017. Protein O-GlcNAcylation: emerging mechanisms and functions. Nat Rev Mol Cell Biol, 18:452–465.

105. Yang Z, Wei X, Ji C, Ren X, Su W, Wang Y, Zhou J, Zhao Z, Zhou P, Zhao K, et al. 2023. OGT/HIF-2alpha axis promotes the progression of clear cell renal cell carcinoma and regulates its sensitivity to ferroptosis. iScience, 26:108148.

106. Yuzwa SA, Macauley MS, Heinonen JE, Shan X, Dennis RJ, He Y, Whitworth GE, Stubbs KA, McEachern EJ, Davies GJ, et al. 2008. A potent mechanism-inspired O-GlcNAcase inhibitor that blocks phosphorylation of tau in vivo. Nat Chem Biol, 4:483–490.

107. Zhang H, Wang Z, Qiao X, Peng N, Wu J, Chen Y, Cheng C. 2024. Unveiling the therapeutic potential of IHMT-337 in glioma treatment: targeting the EZH2-SLC12A5 axis. Mol Med, 30:91.

108. Zhang H, Zhang J, Dong H, Kong Y, Guan Y. 2023. Emerging field: O-GlcNAcylation in ferroptosis. Front Mol Biosci, 10:1203269.

109. Zhang J, Bhuiyan MIH, Zhang T, Karimy JK, Wu Z, Fiesler VM, Zhang J, Huang H, Hasan MN, Skrzypiec AE, et al. 2020. Modulation of brain cation-Cl(-) cotransport via the SPAK kinase inhibitor ZT-1a. Nat Commun, 11:78.

110. Zhang W, Gong M, Zhang W, Mo J, Zhang S, Zhu Z, Wang X, Zhang B, Qian W, Wu Z, et al. 2022. Thiostrepton induces ferroptosis in pancreatic cancer cells through STAT3/GPX4 signalling. Cell Death Dis, 13:630.

111. Zhang X, Sun Y, Ma Y, Gao C, Zhang Y, Yang X, Zhao X, Wang W, Wang L. 2023. Tumor-associated M2 macrophages in the immune microenvironment influence the progression of renal clear cell carcinoma by regulating M2 macrophage-associated genes. Front Oncol, 13:1157861.

112. Zhang Y, Sun C, Ma L, Xiao G, Gu Y, Yu W. 2024. O-GlcNAcylation promotes malignancy and cisplatin resistance of lung cancer by stabilising NRF2. Clin Transl Med, 14:e70037.

113. Zhao CX, Luo CL, Wu XH. 2015. Hypoxia promotes 786-O cells invasiveness and resistance to sorafenib via HIF-2alpha/COX-2. Med Oncol, 32:419.

114. Zhong J, Martinez M, Sengupta S, Lee A, Wu X, Chaerkady R, Chatterjee A, O’Meally RN, Cole RN, Pandey A, et al. 2015. Quantitative phosphoproteomics reveals crosstalk between phosphorylation and O-GlcNAc in the DNA damage response pathway. Proteomics, 15:591–607.

115. Zhou Y, Zhang Q, Zhao Z, Hu X, You Q, Jiang Z. 2024. Targeting kelch-like (KLHL) proteins: achievements, challenges and perspectives. Eur J Med Chem, 269:116270.

116. Zhu G, Murshed A, Li H, Ma J, Zhen N, Ding M, Zhu J, Mao S, Tang X, Liu L, et al. 2021. O-GlcNAcylation enhances sensitivity to RSL3-induced ferroptosis via the YAP/TFRC pathway in liver cancer. Cell Death Discov, 7:83.

117. Zou Y, Palte MJ, Deik AA, Li H, Eaton JK, Wang W, Tseng YY, Deasy R, Kost-Alimova M, Dancik V, et al. 2019. A GPX4-dependent cancer cell state underlies the clear-cell morphology and confers sensitivity to ferroptosis. Nat Commun, 10:1617.

